# Glucocorticoid chrono-pharmacology unveils novel targets for the cardiomyocyte-specific GR-KLF15 axis in cardiac glucose metabolism

**DOI:** 10.1101/2023.12.18.572210

**Authors:** Hima Bindu Durumutla, Ashok Daniel Prabakaran, Fadoua El Abdellaoui Soussi, Olukunle Akinborewa, Hannah Latimer, Kevin McFarland, Kevin Piczer, Cole Werbrich, Mukesh K Jain, Saptarsi M Haldar, Mattia Quattrocelli

## Abstract

Circadian time-of-intake gates the cardioprotective effects of glucocorticoid administration in both healthy and infarcted hearts. The cardiomyocyte-specific glucocorticoid receptor (GR) and its co-factor, Krüppel-like factor (Klf15), play critical roles in maintaining normal heart function in the long-term and serve as pleiotropic regulators of cardiac metabolism. Despite this understanding, the cardiomyocyte-autonomous metabolic targets influenced by the concerted epigenetic action of GR-Klf15 axis remain undefined. Here, we demonstrate the critical roles of the cardiomyocyte-specific GR and Klf15 in orchestrating a circadian-dependent glucose oxidation program within the heart. Combining integrated transcriptomics and epigenomics with cardiomyocyte-specific inducible ablation of GR or Klf15, we identified their synergistic role in the activation of adiponectin receptor expression (*Adipor1*) and the mitochondrial pyruvate complex (*Mpc1/2*), thereby enhancing insulin-stimulated glucose uptake and pyruvate oxidation. Furthermore, in obese diabetic (*db/db*) mice exhibiting insulin resistance and impaired glucose oxidation, light-phase prednisone administration, as opposed to dark-phase prednisone dosing, effectively restored cardiomyocyte glucose oxidation and improved diastolic function towards control-like levels in a sex-independent manner. Collectively, our findings uncover novel cardiomyocyte-autonomous metabolic targets of the GR-Klf15 axis. This study highlights the circadian-dependent cardioprotective effects of glucocorticoids on cardiomyocyte glucose metabolism, providing critical insights into chrono-pharmacological strategies for glucocorticoid therapy in cardiovascular disease.

**GRAPHICAL ABSTRACT:** 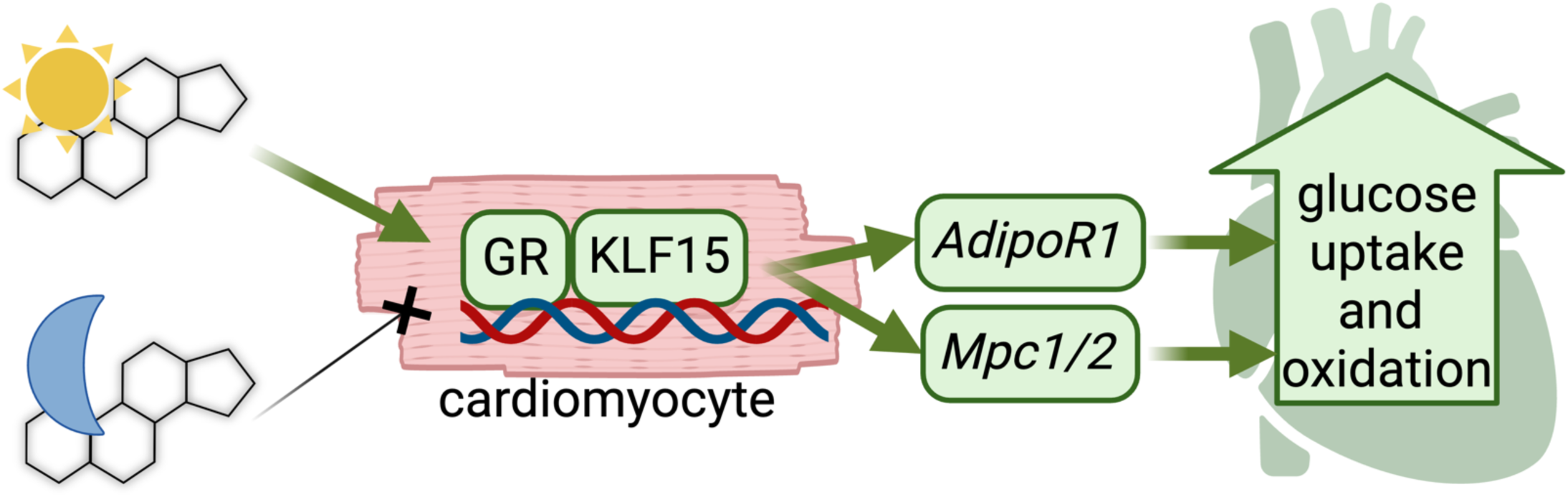

**Brief summary:** Depending on when it is taken during the day, the drug prednisone activates the heart to

## INTRODUCTION

Chronic elevation of glucose levels in diabetes leads to pathological changes in mammalian cells, culminating in cellular and tissue damage, particularly affecting the cardiovascular system. Cardiovascular complications stand as the primary cause of morbidity and mortality among patients with diabetes mellitus (1). Moreover, individuals with metabolic disorders such as obesity and type-2 diabetes (T2D) are predisposed to diabetic cardiomyopathy, a condition of metabolic heart failure (2). T2D imposes metabolic stress on the heart, manifesting in – among others - two major defects: insulin resistance and impaired glucose oxidation. Insulin resistance in glucose uptake constitutes a critical metabolic insult in diabetic hearts and promotes a substrate imbalance that exacerbates diastolic dysfunction, i.e. loss of glucose oxidation with dominating lipid oxidation (3, 4). Moreover, the accumulation of ceramides in the diabetic myocardium further promotes a lipotoxic effect, compromising insulin sensitivity and glucose uptake (5). While previous studies have addressed the deficits in glucose oxidation and insulin sensitivity as single lines of action, comprehensive strategies to manage the metabolic abnormalities in diabetic cardiomyopathy remain elusive. Despite advancements in therapeutics approaches, including SGLT2 inhibitors and GLP1 agonists and/or lifestyle interventions, cardiovascular disease remains a leading cause of morbidity and mortality, accounting for ∼40% of deaths in diabetic patients (6) – a statistic that continues to rise every year. Here we report a completely unanticipated role for the glucocorticoid receptor (GR) and its co-factor Krüppel-like factor 15 (Klf15) in regulating insulin-sensitive glucose uptake and oxidation in heart, addressing a critical clinical need to mitigate cardiovascular morbidity in these patients.

The glucocorticoid receptor (GR) is a potent metabolic regulator that acts as transcription factor when activated by glucocorticoids. The cardiomyocyte-specific GR is important for heart function (7), but the cardiomyocyte-autonomous role of GR in heart metabolism remains unknown. Notably, the glucocorticoid-GR axis is tightly linked to the circadian rhythm (8) and we previously reported that circadian time-of-intake gates the cardioprotective program enabled by the glucocorticoid in heart in a cardiomyocyte-autonomous, clock-dependent way (9). Among its targets, the GR activates its co-factor KLF15 (10), which regulates cardiac metabolism (11) and promotes circadian transcriptional programs in heart (12). However, direct metabolic targets of the GR-KLF15 axis in cardiomyocytes are still remarkably unknown. Here we combine chrono-pharmacology with cardiomyocyte-specific inducible ablation and RNA-seq/ChIP-seq datasets to identify *Adipor1* and *Mpc1/2* as novel critical targets of GR and KLF15 in cardiac glucose utilization. Furthermore, we tested the relevance of these chrono-pharmacology targets for cardiac glucose oxidation and diastolic dysfunction improvements in obese diabetic mice. Our findings challenge the current paradigms in glucocorticoid biology and identify novel druggable mechanisms underlying the interplay between GR and KLF15. These insights may offer potential therapeutic strategies to combat metabolic inflexibility in heart failure.

## RESULTS

### Light-phase dosing gates the glucocorticoid effects on cardiomyocyte glucose metabolism

In our previous study, we reported that the cardioprotective circadian effects of the glucocorticoid prednisone in infarcted murine hearts are gated by the circadian time-of-intake (9). To elucidate the underlying molecular mechanisms, we analyzed RNA-seq datasets from uninjured (sham-operated) and myocardial tissues (GSE186875). Our analysis revealed that intermittent prednisone administration (once-weekly i.p. 1mg/kg for 12 weeks) at Zeitgeber time 0 (ZT0, lights-on) significantly upregulated genes associated with increased glucose import and pyruvate transport pathways compared to vehicle controls. Amongst the genes involved in these pathways, we observed a remarkable upregulation of *Klf15*, *Adipor1* and *Mpc1/2* following ZT0 prednisone stimulation **(Figure 1A)**. Klf15 is a known circadian metabolic regulator of nutrient flux in the heart (12–15), yet its specific targets in conjunction with its co-factor GR (16) remain unknown. *Adipor1* encodes the adiponectin-activated ceramidase that is sufficient to promote insulin-sensitive glucose uptake (17, 18); *Mpc1/2* encode the two subunits of the mitochondrial pyruvate carrier, essential for cardioprotective pyruvate oxidation (19–21). To test whether the steroid-driven transactivation of these genes was dependent on time-of-day, we treated *wild-type* C57BL/6J males with 12-week-long intermittent prednisone treatments at ZT0 versus ZT12, i.e. at start of light-phase vs start of dark-phase **(Supplemental Figure 1A).** Consistent with our RNAseq data, qPCR analysis showed that ZT0 prednisone significantly increased *Klf15, Adipor1, Mpc1 and Mcp2* compared to vehicle and, notably, the drug effect was significantly lower or blunted with ZT12 injections **(Figure 1B)**. While we reported that circadian time of prednisone dosing did not change overall protein levels or nuclear translocation capacity of the GR in heart (9), we were intrigued by the time-of-dosing effects on *Klf15* expression. As KLF15 physically binds the GR as co-activator (16), we tested the extent to which the time-specific gating on *Klf15* upregulation converted in a parallel chrono-pharmacological effect on GR-KLF15 protein-protein interaction. At 4hrs after last drug dose, we found that indeed KLF15 binding to GR was increased by ZT0 prednisone to a higher extent than what shown by ZT12 prednisone **(Figure 1C)**. To assess the functional implications of *Adipor1, Mpc1 and Mcp2* upregulation in cardiac metabolic remodeling, we quantitated myocardial ceramide abundance through mass-spectrophotometry; glucose uptake through 2-deoxyglucose (2DG) assay *in vivo*; glucose oxidation through glucose-fueled respirometry in isolated cardiomyocytes and pyruvate-fueled respiratory control ratio (RCR) (22) in myocardial-derived isolated mitochondria, analyzing data as aggregated **(Figure 1C-F)** and disaggregated by sex **(Figure S1B-E)**. In line with *Adipor1* upregulation, ZT0 but not ZT12 prednisone decreased myocardial ceramides **(Figure 1C)**. With ZT0 prednisone administration, we observed a significant increase in insulin-dependent 2DG uptake **(Figure 1D)**, glucose-fueled respiration, and calculated ATP production in cardiomyocytes **(Figure 1E)**. Consistent with the upregulation of *Mpc1/2*, ZT0 treatment also elevated ADP-stimulated respiration in mitochondria, increasing pyruvate-fueled RCR **(Figure 1F)**. Ceramide clearance and the effects in cardiomyocytes and mitochondria were sex-independent **(Figure S1B-E)**. Regarding the gene expression trends, we sought to gain insight in expression oscillations and time-of-intake effects in heart. We performed a circadian time-course qPCR analysis in myocardial tissues after a single prednisone pulse at ZT0 vs ZT12, to compare circadian trends in the absence of chronic secondary effects. Expression curves revealed oscillations of variable amplitude for *Klf15, Adipor1 and Mpc1/2* in the wild-type myocardium, with the most pronounced circadian mRNA acrophases for *Klf15* in the light-phase and *Mpc1* during the light-dark phase transition **(Figure 1G)**. The transactivation effect by a single prednisone pulse was pronounced and prolonged over the circadian cycle with ZT0 injections, while ZT12 prednisone effects were either transient or non-significant **(Figure 1G)**. Considering the upregulation of *Klf15* appeared central in the circadian-specific prednisone-GR program, we asked whether the cardiomyocyte clock was mediating the time-specific effect. Indeed, we previously showed that the cardiomyocyte-specific BMAL1 was required for the ZT0 prednisone-specific pro-energetic effects in heart (9). Following the ChIP-seq indications of GR-BMAL1 interplay previously garnered in muscle (23), we looked for GR elements (GRE; ACAnnnTGT) and BMAL1 E-Box sites (CACGTG) in the *Klf15* proximal promoter within 300-900bp from each other and found a proximal promoter region with GRE and E-Box at ∼600bp from each other **(Supplemental Figure 1F)**. Through ChIP-qPCR in myocardial tissue at 4hrs after last drug injection, we found that GR occupancy on the GRE was increased by treatment regardless of time-of-intake, while BMAL1 occupancy on the E-Box was increased specifically by ZT0 and not ZT12 prednisone **(Supplemental Figure 1F).** We leveraged the mice with inducible cardiomyocyte-specific BMAL1-KO we previously reported (9) to assess the extent to which BMAL1 was required for the ZT0 prednisone effect on cardiac *Klf15* transactivation. After tamoxifen and washout following prior conditions (9), BMAL1-KO vs BMAL1-WT littermates were treated with 12-week-long ZT0 prednisone or vehicle treatments. In heart, cardiomyocyte BMAL1 ablation blocked the ZT0 prednisone effects on GR occupancy of the Klf15 promoter GRE and on Klf15 mRNA upregulation. The data were further validated by parallel analyses in skeletal muscle, where BMAL1 was not ablated and ZT0 prednisone effects were indeed not blocked **(Supplemental Figure 1G).** Taken together, our findings indicate that circadian time-of-intake gates the prednisone effects on *Klf15, Adipor1 and Mpc1/2* oscillations, as well as ceramide clearance, insulin-sensitive glucose uptake and pyruvate oxidation in the uninjured heart.

**Figure 1.**
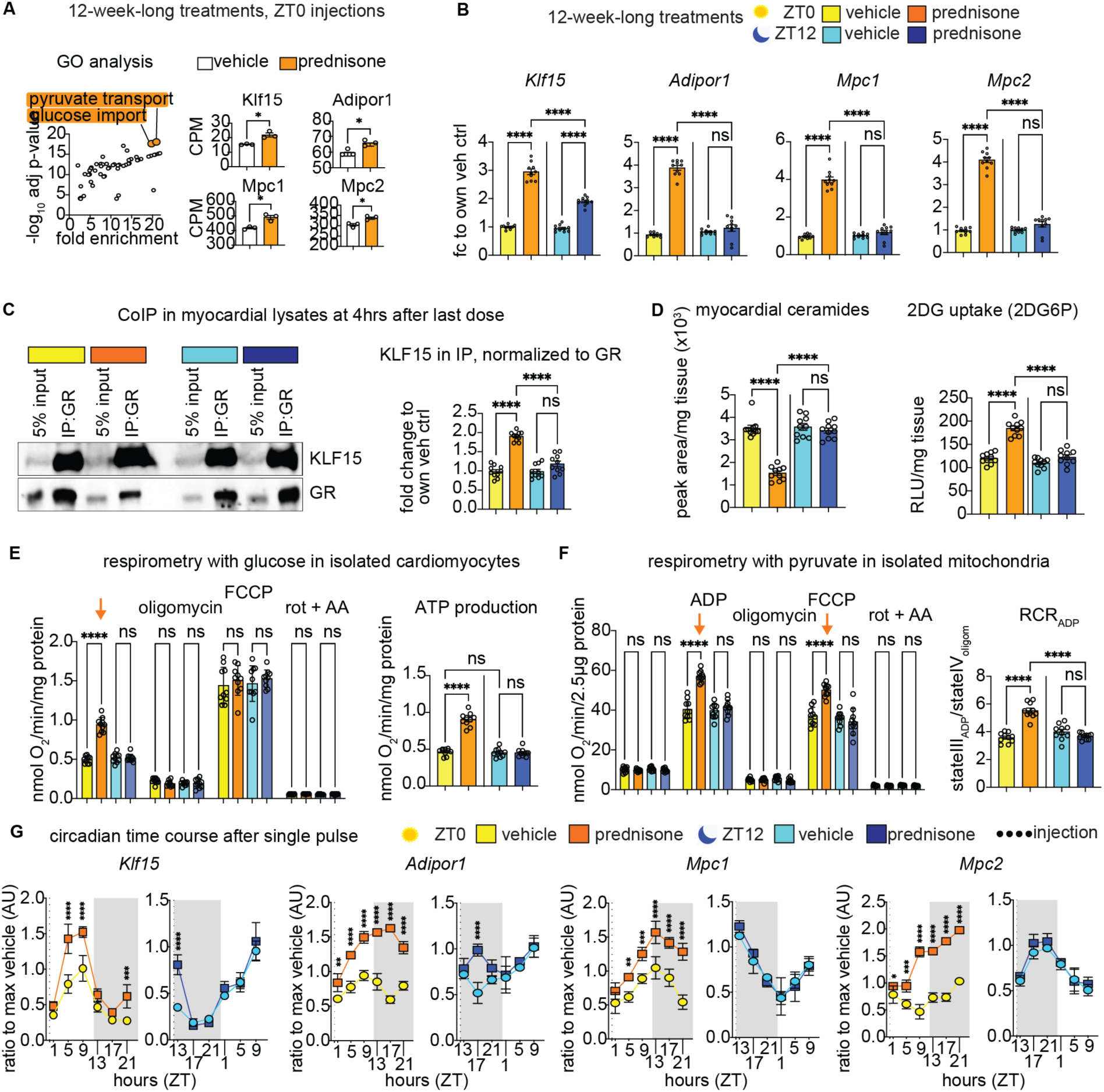
Light-phase glucocorticoid stimulation improves glucose uptake and oxidation in the heart. **(A)** Gene Ontology (GO) analysis revealed that ZT0 intermittent prednisone increased expression of *Klf15*, *Adipor1*, *Mpc1* and *Mpc2* from the enriched pathways of pyruvate transport and glucose import in the wild-type hearts. **(B)** Upregulation of Klf15 by prednisone was higher after ZT0 than ZT12 dosing, and upregulation of *Adipor1*, *Mpc1* and *Mpc2* was specific to ZT0 and blunted by ZT12 dosing. **(C)** CoIPs in heart tissue showed that KLF15 interaction with GR was higher with ZT0 than ZT12 prednisone. **(D)** Consistent with *Adipor1* upregulation, ZT0 prednisone treatment decreased myocardial ceramide levels and increased insulin-dependent 2DG uptake in heart. **(E)** After ZT0 treatment, cardiomyocytes increased basal glucose-fueled respiration ex vivo (arrow), producing more ATP than control or ZT12 treatment. **(F)** Consistent with *Mpc1/2* upregulation, ZT0 treatment increased ADP-stimulated respiration (arrow) and respiratory control ratio with pyruvate. **(G)** Circadian time-course qPCR analyses in myocardial tissues revealed oscillations of variable amplitude for *Klf15*, *Adipor1* and *Mpc1/2* expression. The transactivation effect by a single prednisone pulse was pronounced and prolonged over the circadian cycle with ZT0 injections, while ZT12 prednisone effects were either transient or non-significant. Shown are mean±S.E.M, histograms show also individual mouse values. n=3♂/group in A, n=(5♀+5 ♂)/group in B-G; A: Welch’s t-test; B-H: 2w ANOVA + Sidak; *, P<0.05; **, P<0.01; ***, P<0.001; ****, P<0.0001.

### Generation of ChIP-seq and RNA-seq datasets for the cardiomyocyte-autonomous GR and KLF15

Given the expected upregulation of *Klf15* and activation of GR by prednisone treatment, we tested the role of the GR-Klf15 axis in mediating the upregulation of *Adipor1* and *Mpc1/*2 by light-phase glucocorticoids specifically in cardiomyocytes. For this we sought to garner unbiased evidence of GR and KLF15 epigenomic occupancy and transcriptional effects in heart. To profile epigenomic occupancy of GR and Klf15 in myocardial tissue, we conducted parallel chromatin immunoprecipitation sequencing (ChIP-seq) experiments at 4 hours after a single pulse of ZT0 prednisone administration, adapting the method reported for muscle in our previous study (24). We focused on epigenomic shifts after single drug pulse to avoid secondary effects of chronic drug administration. Notably, due to the lack of ChIP-seq-grade commercial antibodies for Klf15, we performed the immunoprecipitations in myocardial tissue of Klf15-3xFLAG^+/-^ knock-in mice, which were previously reported (25). Our unbiased motif analysis validated the specificity of GR and Klf15 immunoprecipitations, revealing high enrichment for each factor’s canonical binding motifs (glucocorticoid response elements, GREs; KLF response elements, KREs; **Figure 2A)**. The epigenomic activity of both GR and Klf15 was activated by ZT0 prednisone in myocardium, as indicated by increased peak number and signal intensity over canonical sites genome-wide **(Figure 2B)**. However, there were no discernible drug-induced changes in peak localization, which remained highly enriched in promoter-transcriptional start sites (TSS) **(Figure 2B)**.

**Figure 2.**
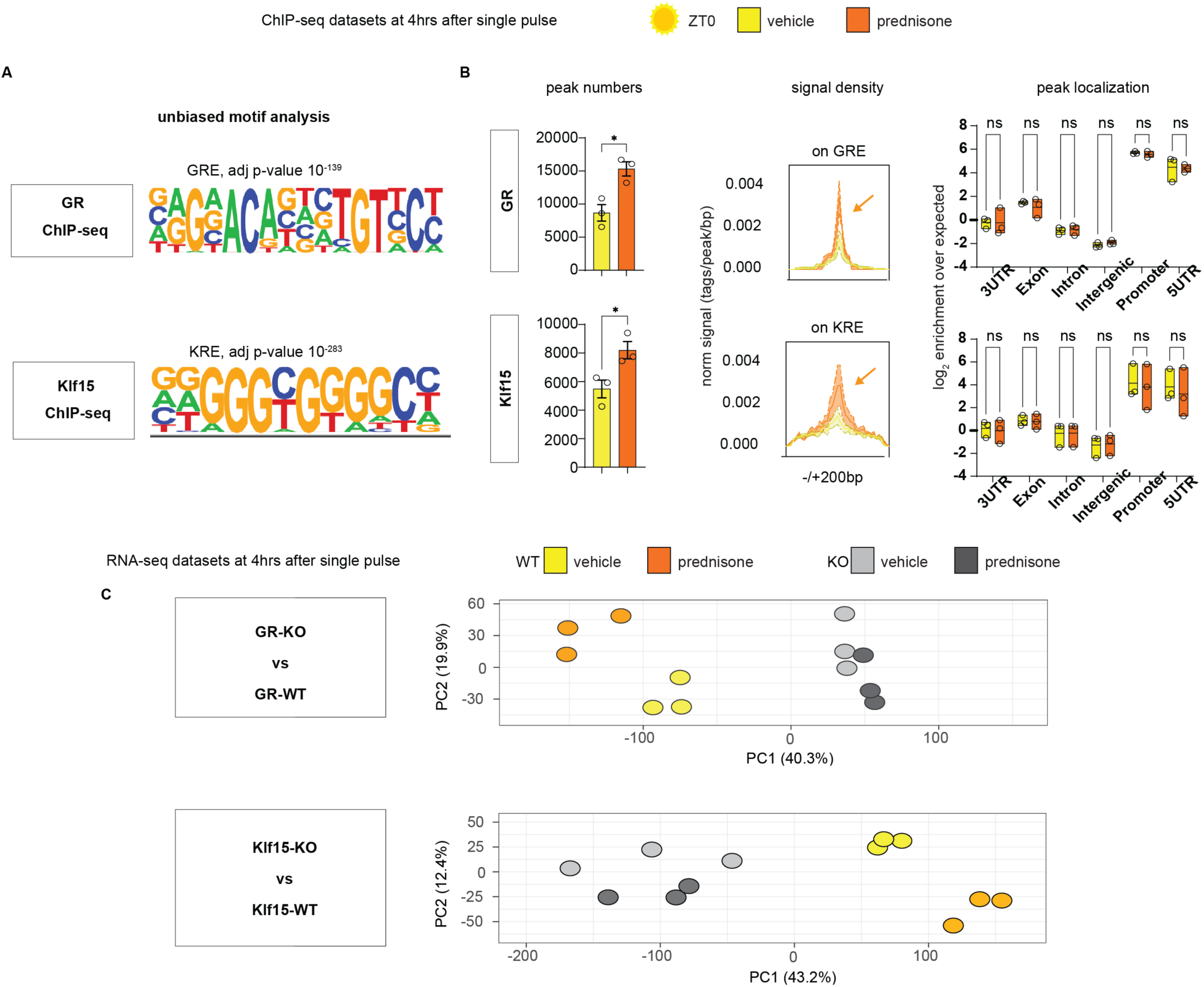
ChIP-seq and RNA-seq datasets reveal non-redundant mediator effect of both cardiomyocyte-specific GR and Klf15 on ZT0 glucocorticoid gene programs in heart. **(A)** Validation of ChIP-seq datasets through unbiased motif analysis. **(B)** Prednisone increased peak numbers and canonical epigenomic peak signal (arrows) for both GR and KLF15 without changes in relative enrichment in genomic localization, which was enriched in promoter and 5’UTRs for both factors. **(C)** PCA analyses show sample clustering according to drug and genotype variables. n=3♂/group; B-D: peak number, Welch’s t-test; peak localization, 2w ANOVA + Sidak; *, P<0.05; **, P<0.01; ***, P<0.001; ****, P<0.0001.

Next, to gain proof of requirement for target gene expression trends, we collected RNA-seq datasets from hearts of cardiomyocyte-specific inducible knockout (*KO*) versus wild-type (*WT*) mice for each factor, using the same tamoxifen and washout protocol **(Figure S2A)**. For GR-KO we used the mice we previously reported (9), comparing hearts from *Myh6-MerCreMer^+/-^;Nr3c1^fl/fl^* (*GR-KO*) versus *Myh6-MerCreMer^+/-^;Nr3c1^wt/wt^* (*GR-WT*) mice. Similarly, for *Klf15-KO* mice we compared *Myh6-MerCreMer^+/-^;Klf15^fl/fl^* (*KLF15-KO*) versus *Myh6-MerCreMer^+/-^;Klf15^wt/wt^* (*KLF15-WT*) **(Supplemental Figure 2B**), using the previously reported *Klf15-floxed* allele (12). As for the ChIP-seq datasets, the RNA-seq datasets were garnered from hearts collected at 4 hours after a single ZT0 prednisone pulse. Principal component analysis (PCA) of the RNA-seq datasets clustered samples based on drug treatment and cardiomyocyte genotype for either *GR* or *Klf15* **(Fig 2C)**, suggesting a non-redundant requirement for both factors in the transcriptional program enabled by ZT0 prednisone in heart. We therefore generated datasets to unbiasedly probe the extent to which the cardiomyocyte-specific GR-KLF15 axis regulates *Adipor1 and Mpc1/2* expression in heart.

### Cardiomyocyte GR and Klf15 are independently required for the ZT0 prednisone effect on Adipor1 and Mpc1/2 transactivation, insulin-driven glucose uptake and pyruvate oxidation

We first tested the extent to which *Adipor1* was a direct transactivation target of GR and Klf15 in heart downstream of ZT0 prednisone. ChIP-seq peak tracks revealed steroid-sensitive peaks on the *Adipor1* proximal promoter for both GR and Klf15. RNA-seq data showed that the inducible cardiomyocyte-specific ablation of either GR or Klf15 blocked the drug-induced upregulation of *Adipor1*, indicating a non-redundant requirement for both factors in a cardiomyocyte-autonomous fashion **(Figure 3A)**. Based on the motifs unveiled by the unbiased motif analysis of ChIP-seq datasets (ACAnnnTGT for GRE; GGGCGGGG for KRE), we found GRE and KRE motifs in the proximal promoter of *Adipor1* and validated the ChIP-seq and RNA-seq indications through ChIP-qPCRs and qPCRs **(Supplemental Figure 2C)**. In accordance with the trends in *Adipor1* transactivation, cardiomyocyte-specific inducible ablation of either GR or Klf15 blocked the chronic ZT0 prednisone effects on ceramide reduction and 2DG uptake in heart. The transcriptional and metabolic effects of treatment appeared durable and extended to not only the light-phase, but also the active phase of treated mice **(Figure 3B; Supplemental Figure 2C)**. Similarly, we found steroid-sensitive peaks on the proximal promoters of *Mpc1* and *Mpc2*, with ZT0 prednisone transactivation of both genes blocked by cardiomyocyte-specific ablation of either GR or Klf15 **(Figure 3C)**. We found GRE and KRE motifs in the 5’ UTRs of both *Mpc1* and *Mpc2* and validated GR-KLF15 occupancy and gene upregulation through ChIP-qPCRs and qPCRs **(Supplemental Figure 2C)**. In accordance with the trends in *Mpc1/2* transactivation, the chronic treatment-driven gain in pyruvate oxidation was also blocked by the inducible gene and the KLF15-dependent chronic treatment effects were applicable to both active and rest phase of mice (**Figure 3D; Supplemental Figure 2D**). Taken together, these findings demonstrate that the ZT0-specific prednisone stimulation of glucose uptake and oxidation in heart is mediated by the cardiomyocyte-autonomous epigenetic activity of both GR and Klf15.

**Figure 3.**
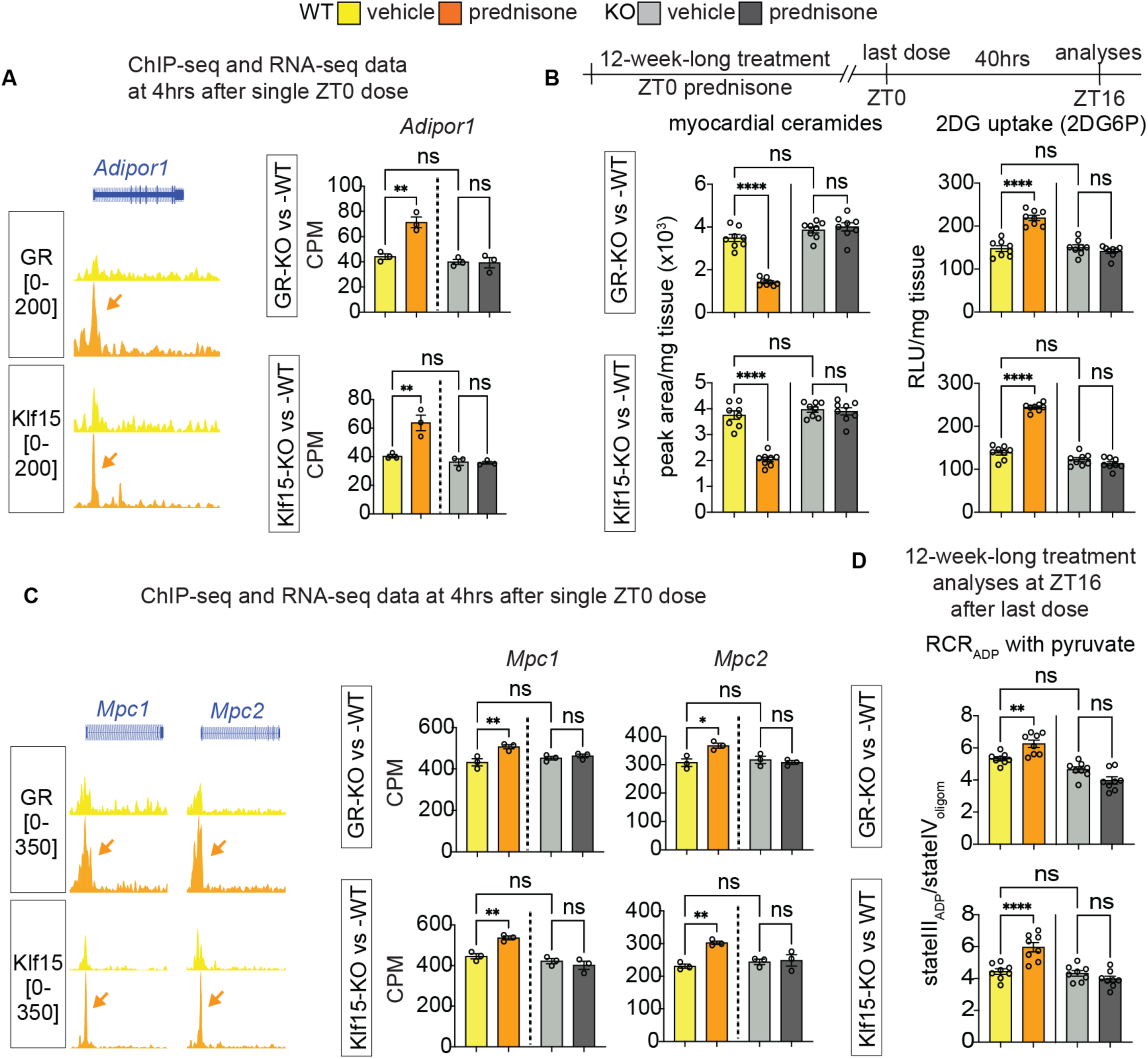
Cardiomyocyte GR and KLF15 are required for the ZT0 prednisone effects on gene transactivation, insulin sensitization and pyruvate oxidation in heart. **(A)** GR and KLF15 showed steroid-sensitive peaks on *Adipor1* TSS (arrows). Drug-induced upregulation was blunted by inducible cardiomyocyte-specific ablation of either GR or KLF15. **(B)** Transcriptional/metabolic effects of treatment appeared durable beyond the rest phase as GR and KLF15 were both required for the treatment effects on myocardial ceramide content reduction and insulin-stimulated glucose uptake increment in heart in the active phase (ZT16). **(C-D)** Similarly, GR and KLF15 were required for the drug effect on *Mpc1/2* transactivation, and for the chronic treatment effects on pyruvate oxidation in heart in the active phase. Shown are mean±S.E.M, histograms show also individual mouse values. n=3♂/group in A, C; n=8♂/group in B, D; 2w ANOVA + Sidak; *, P<0.05; **, P<0.01; ***, P<0.001; ****, P<0.0001.

We then assessed heart function through echocardiography in our cohorts at start and end of 12-week-long ZT0 prednisone treatments. We primarily focused on fractional shortening, stroke volume and heart weight/tibia length, as those parameters were previously shown to be impacted by both GR and KLF15 constitutive (post-natal, not inducible) ablations in heart (26, 27). In our own mice after inducible ablation and in the absence of prednisone, KO for either factor significantly depressed fractional shortening and stroke volume, while increasing heart mass, when compared to WT controls. In WT uninjured hearts, ZT0 prednisone treatment increased stroke volume and fractional shortening, with no significant trends in normalized heart weight. As expected, treatment had no sizable effects in KO hearts **(Figure 4)**. In line with the trends in fractional shortening and stroke volume, KO for either factor enlarged the systolic left ventricle diameter (LVIDs) and induced non-significant declines in ejection fraction and gains in diastolic left ventricle diameter (LVIDd). In WT hearts, treatment induced a small significant reduction of LVIDs, a non-significant trend in increased ejection fraction and no appreciable changes in LVIDd. In KO hearts, treatment had no sizable effects. Heart rate during the echocardiographic measurements, i.e. under anesthesia, was not changed by either KO or treatment **(Supplemental Figure 3)**. Therefore, inducible ablation of either GR or KLF15 after normal growth recapitulated the constitutive post-natal ablation effects on uninjured heart, blocking the treatment-driven gains in stroke volume.

**Figure 4.**
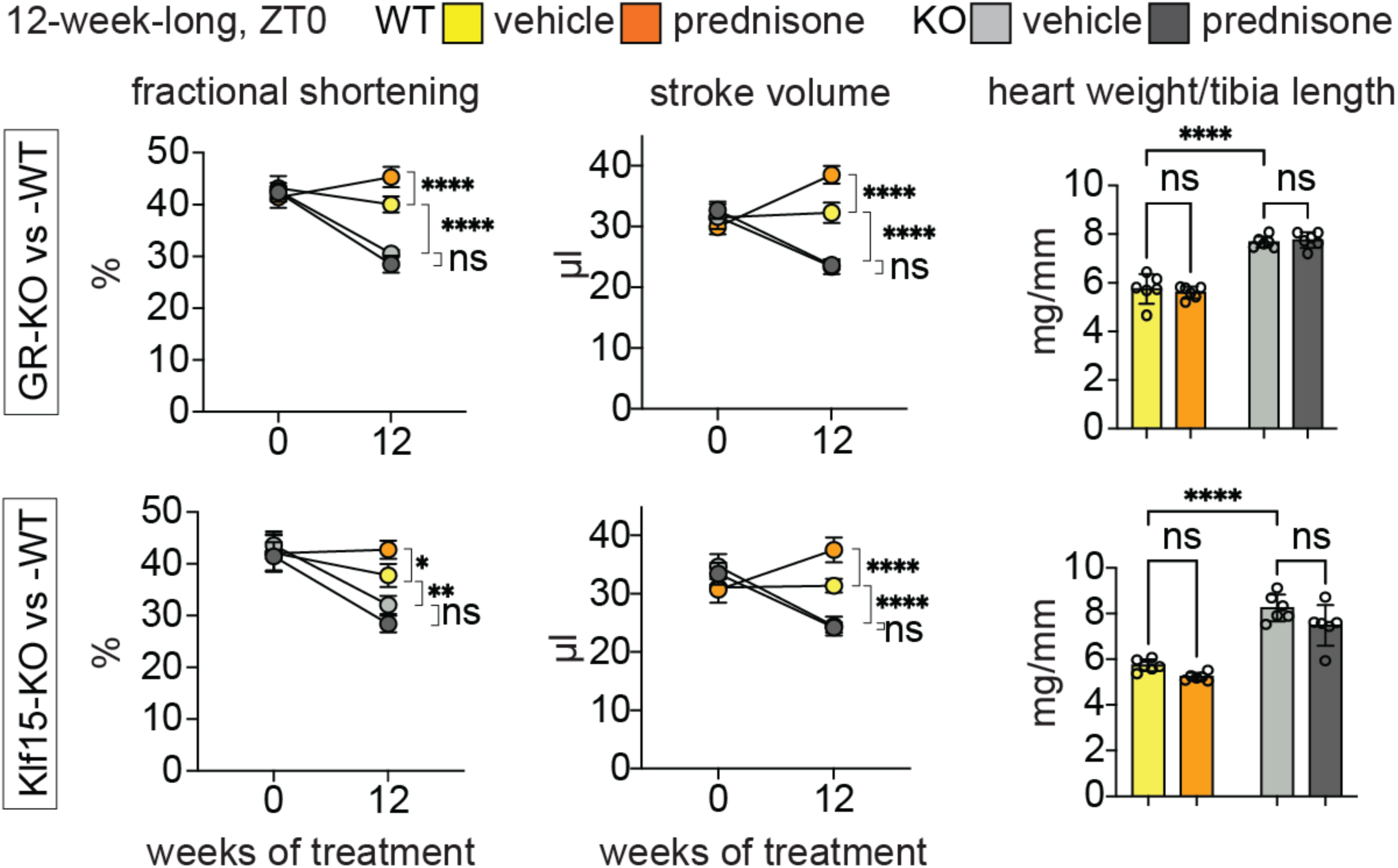
Functional cardiac assessments for chronic ZT0 prednisone effects on uninjured hearts after cardiomyocyte gene ablation. Either GR-KO or KLF15-KO induced significant losses in fractional shortening, stroke volume and gains in heart weight/tibia length compared to WT. In WT uninjured hearts, treatment increased stroke volume and fractional shortening, with no significant trends in heart weight. As expected, treatment had no sizable effects in KO hearts. Shown are mean±S.E.M, histograms show also individual mouse values. N=6♂/group; curves, 3w ANOVA + Sidak; histograms, 2w ANOVA + Sidak; *, P<0.05; **, P<0.01; ***, P<0.001; ****, P<0.0001.

### Light-phase glucocorticoid intermittence rescues glucose oxidation and diastolic dysfunction in diabetic hearts

We hypothesized that combining the ZT0 prednisone effects on insulin sensitization and pyruvate oxidation could serve as a “two-hit” approach to enhance a functional glucose oxidation program in the diabetic heart. Given that insulin resistance and impaired glucose oxidation are key metabolic dysfunctions in T2D diabetic cardiomyopathy (3, 4), we aimed to evaluate the relevance of this ZT0 prednisone program in a diabetic cardiomyopathy model. To address this, we employed 6-month-old *db/db* obese diabetic mice (**Figure 5A**) (28, 29) and their *db/+* littermates as non-obese, non-diabetic parallel control cohort. At baseline, we reconfirmed previously reported defects in diastolic function (mitral valve flow impairment and increased E/e’ (29)), systolic function (decreased stroke volume (30, 31)) and cardiac hypertrophy (increased heart mass to tibia length (32)) in *db/db* mice vs *db/+* controls **(Figure 5B)**. Importantly, these parameters were rescuable in *db/db* mice through experimental interventions re-activating glucose uptake and oxidation (30–33). After treatment with intermittent prednisone at ZT0 versus ZT12 for 12 weeks, ZT0 prednisone alleviated diastolic dysfunction, stroke volume impairment and cardiac hypertrophy in *db/db* mice towards control *db/+* levels, while the treatment effects were blocked by ZT12 injections **(Figure 5C-E)**. Only with ZT0 dosing did treatment increase glucose uptake in myocardial tissue, measured as 2DG uptake, in both *db/+* and *db/db* hearts **(Figure 5F)**. Also, compared to isochronic vehicle, ZT0 but not ZT12 prednisone treatment increased glucose-fueled basal respiration and ATP production in *db/db* cardiomyocytes towards control-like control *db/+* levels (**Figure 5F**). Notably, none of these parameters showed significant sexual dimorphism in their effects (**Supplemental Fig 4A**). Treatments had no sizable effects on body weight beyond the expected genotype-dependent increase **(Supplemental Figure 4B)**. Furthermore, we checked blood pressure through tail cuff measurements on the day after the last injection, considering the hypertensive phenotype of db/db mice (34) and the hypertensive potential of chronic glucocorticoid treatments (35). We conducted the tail cuff measurements in the active phase (ZT16) to avoid potential confounding dips induced by sleep or rest. None of the treatments had sizable effects on blood pressure or heart rate during blood pressure measurements beyond the expected db/db-related mild hypertension in both systole and diastole (**Supplemental Fig 4C**). In light of the ZT0-specific effects of prednisone on KLF15-GR interaction in WT non-diabetic hearts, we tested the extent to which the chronic ZT0 vs ZT12 regimens changed GR or KLF15 nuclear translocation in db/db hearts. We performed WB for GR and KLF15 in db/db myocardial lysate fractions at 30min after the last injection of ZT0 vs ZT12 prednisone at the end of the 12-week-long treatment. We found that time-of-intake did not sizably change the rate of GR nuclear translocation, while it did impact KLF15 translocation, which was specifically increased by ZT0 prednisone **(Supplemental Figure 5A)**. Indeed, in line with the model of GR-KLF15 transactivation of AdipoR1 and MPC1/2, we found that at 24hrs after last drug injection those proteins were increased by ZT0 prednisone – but not ZT12 prednisone – compared to vehicle in db/db hearts **(Supplemental Fig 5B)**. We also found analogous trends in the glucose transporters GLUT1 and GLUT4 **(Supplemental Fig 5B)**, which were convergent with the glucose uptake reactivation in those hearts and consistent with prior seminal findings with KLF15 overexpression (36). Finally, we sought to validate that the ZT0 prednisone effects in db/db hearts were dependent on GR and KLF15. To that goal, we used MyoAAV-serotyped vectors containing either a scramble or anti-*Nr3c1* (MyoAAV-antiGR) or anti-*Klf15* (MyoAAV-antiKLF15) shRNA under the U6 promoter. The MyoAAV serotype maximizes tropism of transduction for AAV vectors in vivo to striated muscles (37). To maximize knockdown, three different vectors each with a different shRNA were combined per gene knockdown. At 6 months of age, male db/db mice were injected r.o. with 6×10^12^ vg/mouse MyoAAV-scramble or 10^12^ vg/mouse/vector MyoAAV-antiGR (3 vectors) + MyoAAV-antiKLF15 (3 vectors) immediately before starting the 12-week-long ZT0 prednisone vs vehicle intermittent regimens. At endpoint, WB in myocardial tissue confirmed GR and KLF15 knockdown (approximately ∼80% protein reduction) and showed that the knockdown vectors blocked or blunted the ZT0 prednisone effects on upregulation of Adipor1, Mpc1/2 protein levels in db/db hearts **(Figure 6A)**. Accordingly, the combinatorial GR+KLF15 knockdown blocked the treatment effects on diastolic dysfunction (E/e’), stroke volume, cardiac hypertrophy (heart weight/tibia length) and insulin-driven glucose uptake (2DG6P; **Figure 6B-C**). Moreover, treatment increased glucose-fueled respiration (oxygen consumption rate, OCR) in permeabilized cardiac biopsies and pyruvate-fueled respiration (respiratory control ratio, RCR) in isolated myocardial mitochondria in scramble- but not knockdown-transduced hearts **(Figure 6D)**. Taken together, these data indicate the potential for prednisone chrono-pharmacology to reactivate glucose oxidation and alleviate dysfunction in obese diabetic hearts by engaging the GR-KLF15 interplay in heart in vivo.

**Figure 5.**
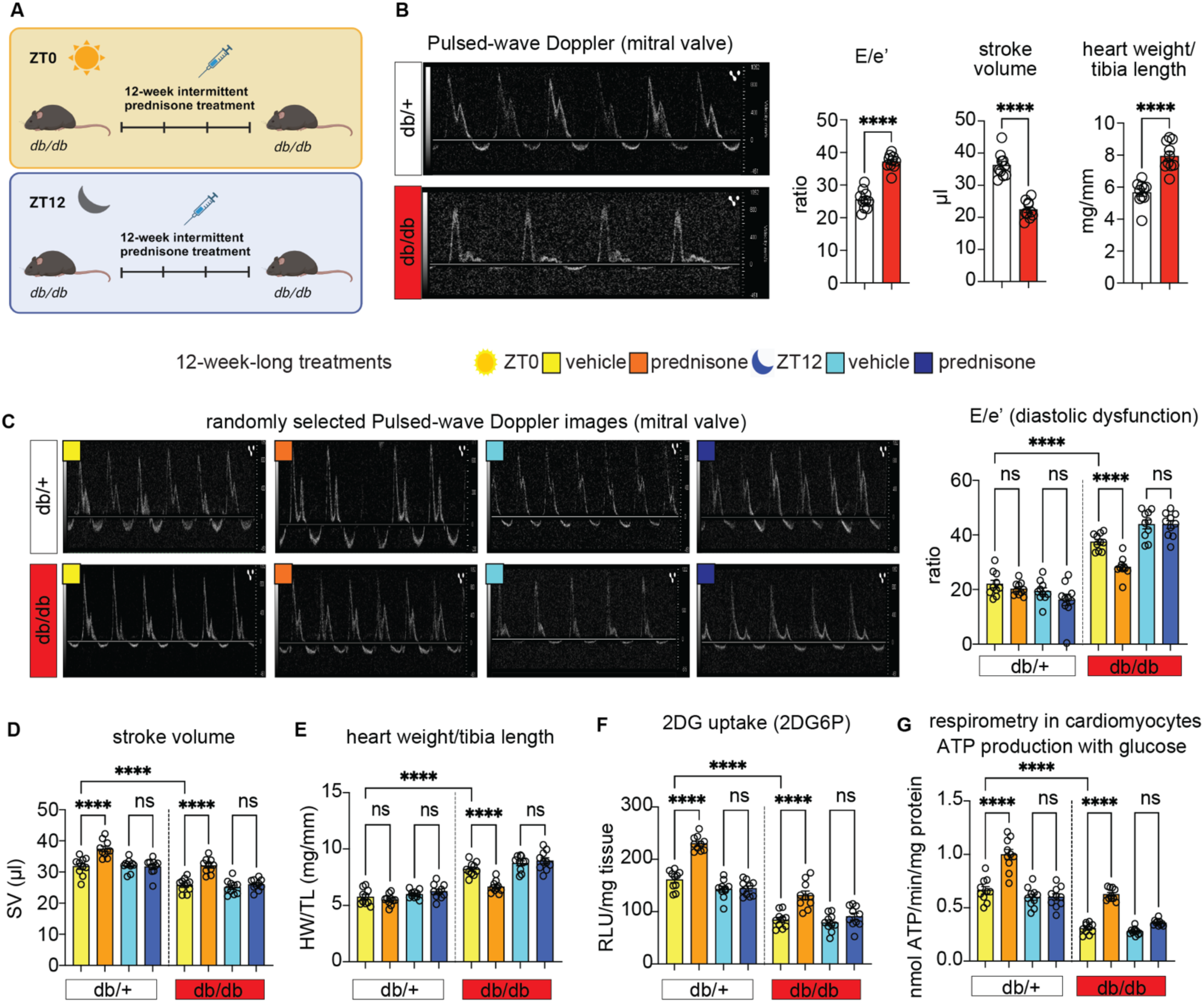
ZT0 glucocorticoid treatment rescues glucose oxidation and diastolic function in db/db mice. **(A)** Schematic showing the 12-week-long prednisone treatment of db/db and their littermate controls (db/+) at ZT0 and ZT12. **(B)** Representative images of pulse wave doppler echocardiography for mitral valve flow, and quantitation of baseline defects in E/e’, stroke volume and heart mass in db/db mice. **(C)** ZT0 treatment decreased E/e’ in db/db hearts to db/+-like levels, while ZT12 treatment had no sizable effects. **(D)** ZT0 but not ZT12 treatment increased stroke volume in both db/db and db/+ mice. **(E)** Cardiac hypertrophy was alleviated in db/db hearts by ZT0 but not ZT12 treatment. **(F)** Treatment increased glucose uptake in myocardial tissue, measured as 2DG uptake, in both db/+ and db/db hearts only with ZT0 dosing. **(G)** Treatment increased glucose-fueled ATP production calculated from Seahorse curves in isolated cardiomyocytes in both db/+ and db/db mice according to time of intake (ZT0 but not ZT12), increasing db/db values to control-like levels. Shown are mean±S.E.M, histograms show also individual mouse values. n=(5♀+5 ♂)/group; B, Welch’s t-test; C-G, 3w ANOVA + Sidak; *, P<0.05; **, P<0.01; ***, P<0.001; ****, P<0.0001.

**Figure 6.**
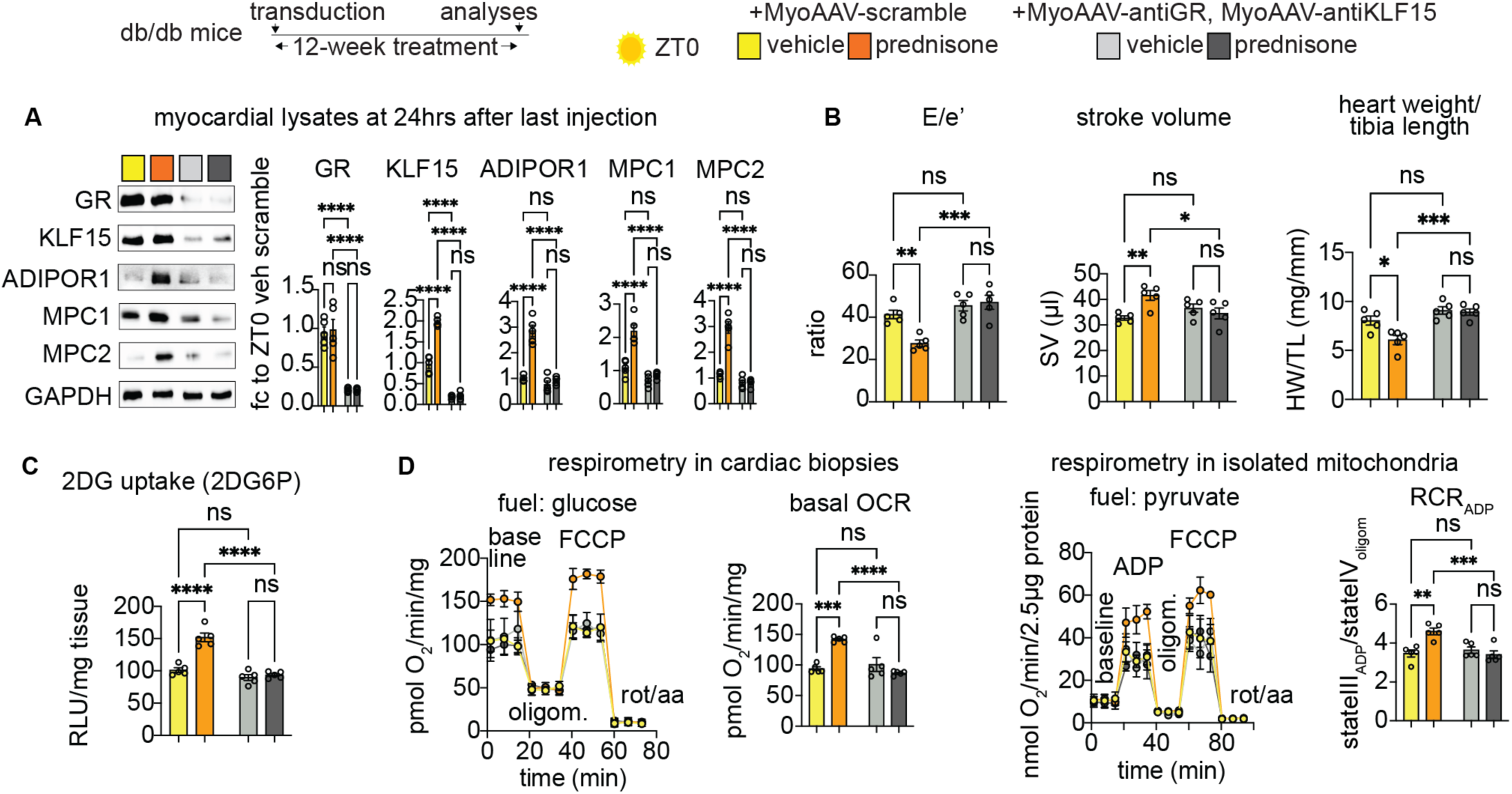
Knock-down of GR and KLF15 in db/db hearts in vivo blocks the ZT0 prednisone effects. **(A)** Representative WB showing knock-down of GR and KLF15 at 12-weeks after transduction and ZT0 prednisone regimen start, as well as blunting of the treatment effect on Adipor1, Mpc1 and Mpc2 protein level upregulation in hearts of db/db mice transduced with the knock-down vectors compared to mice transduced with scramble vectors. **(B-C)** Knock-down MyoAAV combination blocked the treatment effects on diastolic dysfunction (E/e’), stroke volume, cardiac hypertrophy (heart weight/tibia length) and insulin-driven glucose uptake (2DG6P). **(D)** Treatment increased glucose-fueled respiration (oxygen consumption rate, OCR) in permeabilized cardiac biopsies and pyruvate-fueled respiration (respiratory control ratio, RCR) in isolated myocardial mitochondria in scramble- but not knockdown-transduced hearts. Shown are mean±S.E.M, histograms show also individual mouse values. n=5 ♂/group; 2w ANOVA + Sidak; *, P<0.05; **, P<0.01; ***, P<0.001; ****, P<0.0001.

## DISCUSSION

The GR plays a pivotal role in metabolic regulation, acting as a transcription factor when activated by glucocorticoids. While the importance of cardiomyocyte-specific GR in heart function is recognized (7), its autonomous role in heart metabolism remains unclear. Our previous work has elucidated two critical time dimensions influencing the outcomes of GR activation: i) circadian time-of-intake and ii) chronic frequency-of-intake. Light-phase dosing of prednisone in mice has shown benefits in heart function compared to dark-phase dosing (9). Intermittent once-weekly dosing reversed the dysmetabolic effects of chronic daily dosing and improved exercise tolerance (38). In obese diabetic mice, endogenous glucocorticoids are elevated in response to metabolic stress and yet they maintain circadian oscillations, albeit dimmed (39). This will be an important point to elucidate for the translation of our findings into pharmacological strategies for diabetic cardiomyopathy, although pilot clinical data with prednisone chrono-pharmacology in patients with muscular dystrophies showed promising improvements in lean mass and exercise tolerance (40).

Despite the known regulation of GR activity by the circadian clock (41, 42), the specific targets of glucocorticoid chrono-pharmacology for functional cardiac metabolic remodeling remain poorly elucidated. Our findings from this study discover one such mechanism: the cardiomyocyte-autonomous requirement for the GR-KLF15 axis in facilitating light-phase-specific glucocorticoid transactivation of Adipor1 and the MPC complex genes in the heart. Adipor1, an adiponectin-responsive ceramidase (43) is generally linked to pro-metabolic effects and plays a crucial role in insulin-sensitive glucose uptake by degrading ceramides (18, 44). Ceramides accumulate in heart with obesity and/or T2D, promoting cardiac insulin resistance and the related cardiac dysfunction (45). It is important to note that overt, unbalanced reactivation of glucose uptake in the insulin resistant heart causes unwanted side effects, as shown by cardiomyocyte GLUT4 overexpression in diabetic hearts (46). In that perspective, our findings with the simultaneous upregulation of both Adipor1 and the MPC complex could offer a model to reactivate both glucose uptake and downstream oxidation in a balanced fashion.

MPC expression is downregulated in failing human hearts (47) and transcriptional MPC abundance directly promotes cardioprotective oxidation of pyruvate and its precursor glucose in the myocardium (19, 20, 47). Accordingly, MPC activity is decreased in the diabetic heart too (48). It was remarkable to observe a strong concerted epigenetic program by the cardiomyocyte-specific GR and Klf15 on both subunit genes for the MPC, which indeed resulted in pyruvate oxidation trends in line with increased glucose oxidation.

The variable of circadian time of drug intake inevitably questions the time gating on not only basic mechanisms but also chronic effects with respect to the circadian cycle. Here, the ZT0-prednisone-gated interaction between GR and KLF15 stimulated cardiomyocyte-autonomous transcriptional/metabolic effects that extended to both rest- and active-phases of mice after chronic treatment. In other words, a repetitive series of circadian-specific drug stimulations supported a virtuous program that improved glucose handling in cardiomyocytes throughout the circadian cycle. This is an emerging aspect in various contexts of chrono-pharmacology, including the heterogenous landscape of chrono-immunology for cardiovascular health (49). Particularly in the context of diabetic cardiomyopathy, these findings open the question for future studies of which chrono-regimens can be complemented to concertedly leverage the endogenous rhythms of all the critical cell compartments of the heart, including cardiomyocytes, fibroblasts, endothelium and immune cells.

### Limitations of the study

While sufficiently powered for overall trends with chrono-treatments and gene ablation, our cohort numbers were limited and likely underpowered for omics analyses and sex-specific differences. The limited numbers used for RNA-seq and ChIP-seq were mitigated by dataset overlay and further validation through two different cardiomyocyte-specific inducible KO models. The limited numbers of mice in our sex-disaggregated analyses probably limited the detection of sexual dimorphism, which will have to be addressed for the pathways in play here with appropriate cohort numbers and hormonal manipulations in future studies. Despite the mechanistic evidence offered by cardiomyocyte-specific GR and KLF15 inducible ablation in non-diabetic conditions and by the in vivo knockdown in db/db hearts, in this study we did not directly address the requirement for the effects on insulin sensitivity and glucose oxidation per se in the metabolic remodeling of the obese diabetic heart. We believe this is beyond the scope of this investigation.

### Conclusions and overall impact

In summary, our study challenges the current paradigm of glucocorticoid cardiotoxicity by identifying a specific mechanism that mediates the circadian-gated glucocorticoid benefit on cardiomyocyte glucose oxidation, with functional implications for diabetic cardiomyopathy.

## Acknowledgements

Mass-spec analyses were performed at the Cincinnati Children’s Mass-Spec (Clinical and Biomedical) Facility (RRID: SCR_022638), with critical assistance by Drs. Setchell and Zhao. Next-gen sequencing was performed at the Cincinnati Children’s DNA Sequencing and Genotyping Facility (RRID: SCR_022630), with critical assistance by David Fletcher, Keely Icardi, Julia Flynn, and Taliesin Lenhart.

## Supplementary Figures

**Supplementary Figure 1.**
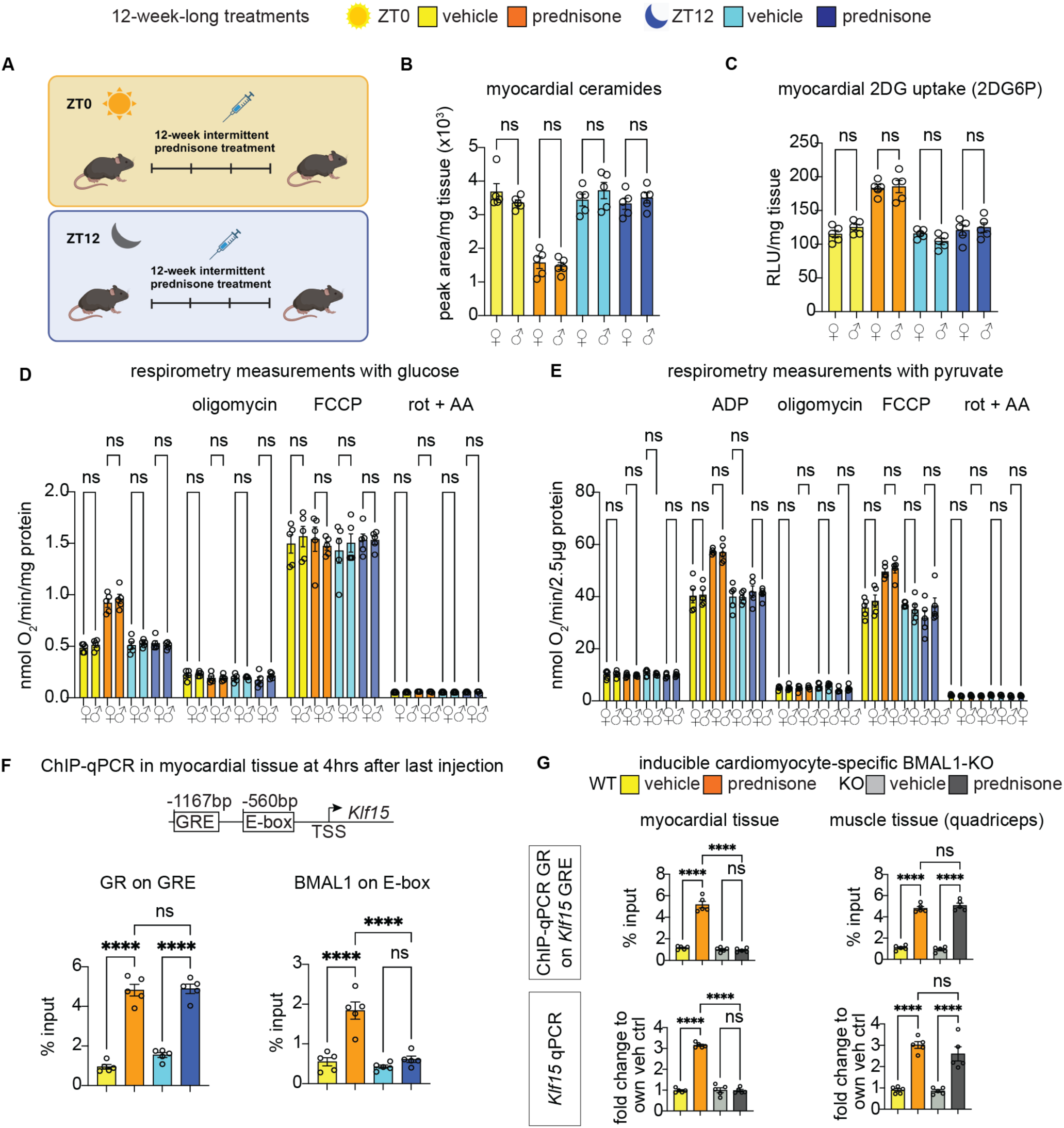
Additional analyses related to. Figure 1**. (A)** Graphical experimental design for wild-type mice. **(B)** Ceramide levels were decreased upon treatment with prednisone at ZT0, but not at ZT12. This effect was sex-independent. **(C)** Increased insulin-dependent 2DG uptake in heart showed no sexual dimorphism. **(D)** Although the cardiomyocytes showed an increase in basal glucose-fueled respiration ex vivo, no differences between the two sexes were observed. **(E)** ZT0 treatment increased ADP-stimulated respiration and respiratory control ratio with pyruvate. These effects were sex-independent. **(F)** While GR occupancy of *Klf15* promoter was increased by treatment independent from time-of-intake, ZT0 but not ZT12 treatment increased BMAL1 recruitment. **(G)** Ablation of BMAL1 in adult cardiomyocytes in vivo blocked the ZT0 prednisone-driven effects on GR occupancy of *Klf15* promoter and *Klf15* mRNA upregulation in heart (BMAL1 ablation) but not in muscle (no BMAL1 ablation). Shown are mean±S.E.M, histograms show also individual mouse values. n=(5♀+5♂)/group B-E, 5♂/group in F-G; 2w ANOVA+ Sidak; ns= non-significant. *, P<0.05; **, P<0.01; ***, P<0.001;

**Supplementary Figure 2.**
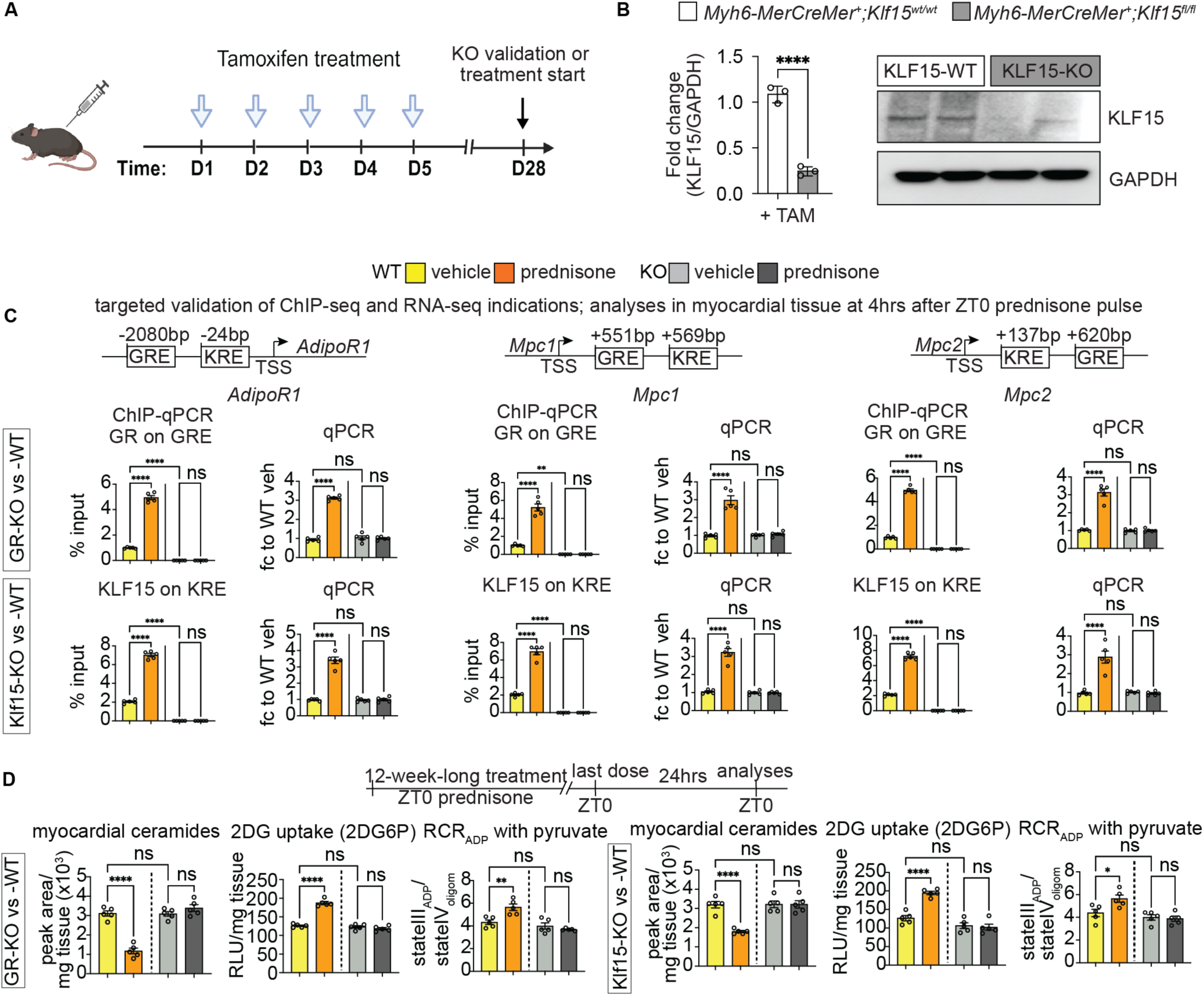
Tamoxifen regimen, Klf15-KO validation and light-phase effects of chronic treatments in GR-KO and KLF15-KO hearts. **(A)** Schematic representing the generation and washout of tamoxifen-mediated inducible knockout mice for GR or KLF15. **(B)** Validation of KLF15 ablation in heart through WB. **(C)** Targeted validation through ChIP-qPCRs and qPCRs of ChIP-seq and RNA-seq indications of GR and KLF15 transactivation of *AdipoR1*, *Mpc1* and *Mpc2* in heart after ZT0 prednisone. **(D)** Related to Figure 3B-D. GR and KLF15 were both required for treatment effects on myocardial ceramides, glucose uptake and pyruvate oxidation at 24hrs from last ZT0 prednisone dose, i.e. in the rest phase. Shown are mean±S.E.M, histograms show also individual mouse values. B, n=3/group, Welch’s t-test; C, n=5♂/group, 2w ANOVA + Sidak; *, P<0.05; **, P<0.01; ***, P<0.001; ****, P<0.0001.

**Supplementary Figure 3.**
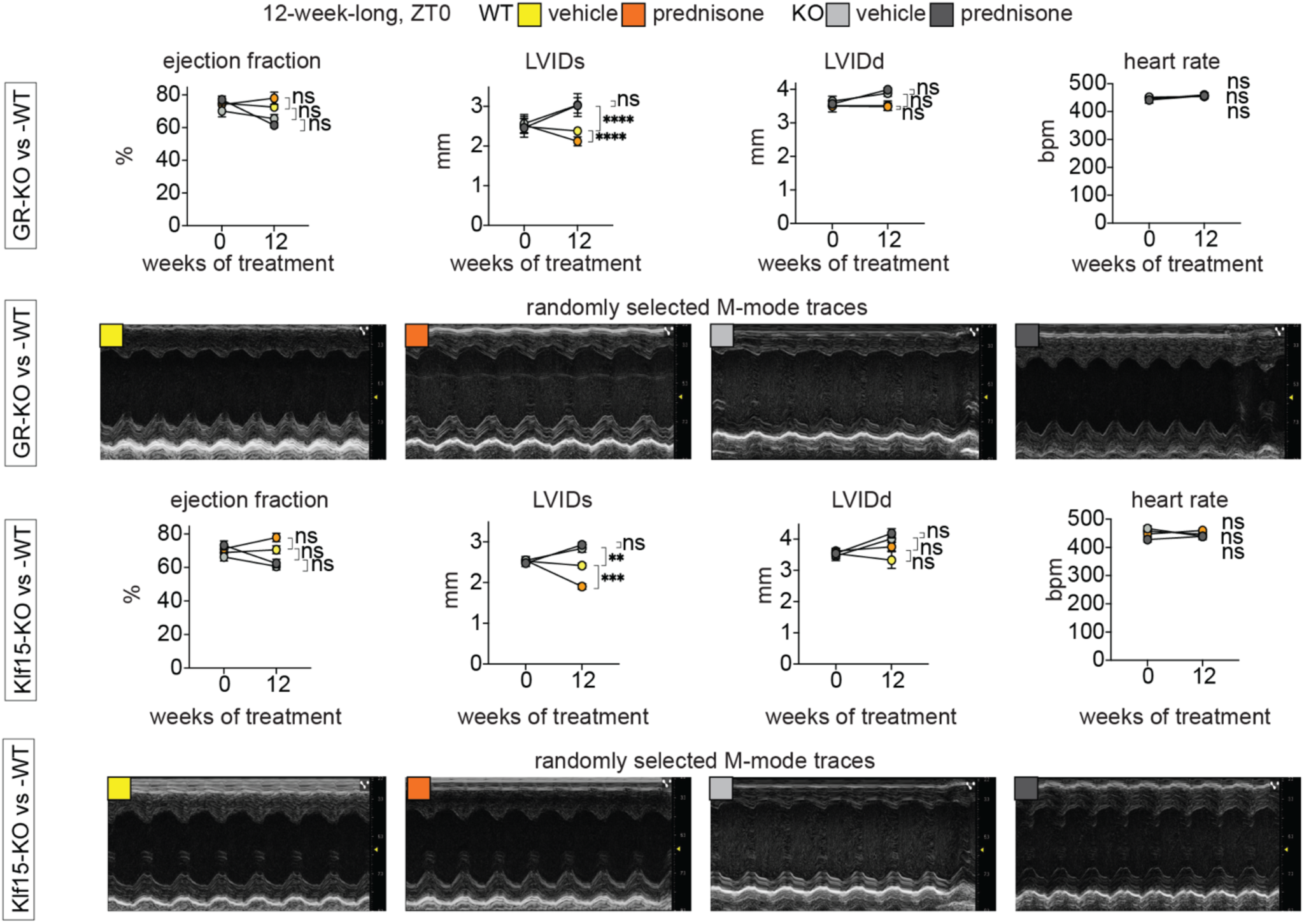
Additional echocardiographic measurements related to. Figure 4. In line with the trends in fractional shortening and strole volume (Figure 4), inducible ablation of either GR or KLF15 in cardiomyocytes enlarged the systolic left ventricle diameter (LVIDs) and induced non-significant declines in ejection fraction and gains in diastolic left ventricle diameter (LVIDd). In WT hearts, treatment induced a small significant reduction of LVIDs, a non-significant trend in increased ejection fraction and no appreciable changes in LVIDd. In KO hearts, treatment had no sizable effects. Heart rate during the echocardiographic measurements, i.e. under anesthesia, was not changed by either KO or treatment. Shown are mean±S.E.M, histograms show also individual mouse values. N=6♂/group; curves, 3w ANOVA + Sidak; histograms, 2w ANOVA + Sidak; *, P<0.05; **, P<0.01; ***, P<0.001; ****, P<0.0001.

**Supplementary Figure 4.**
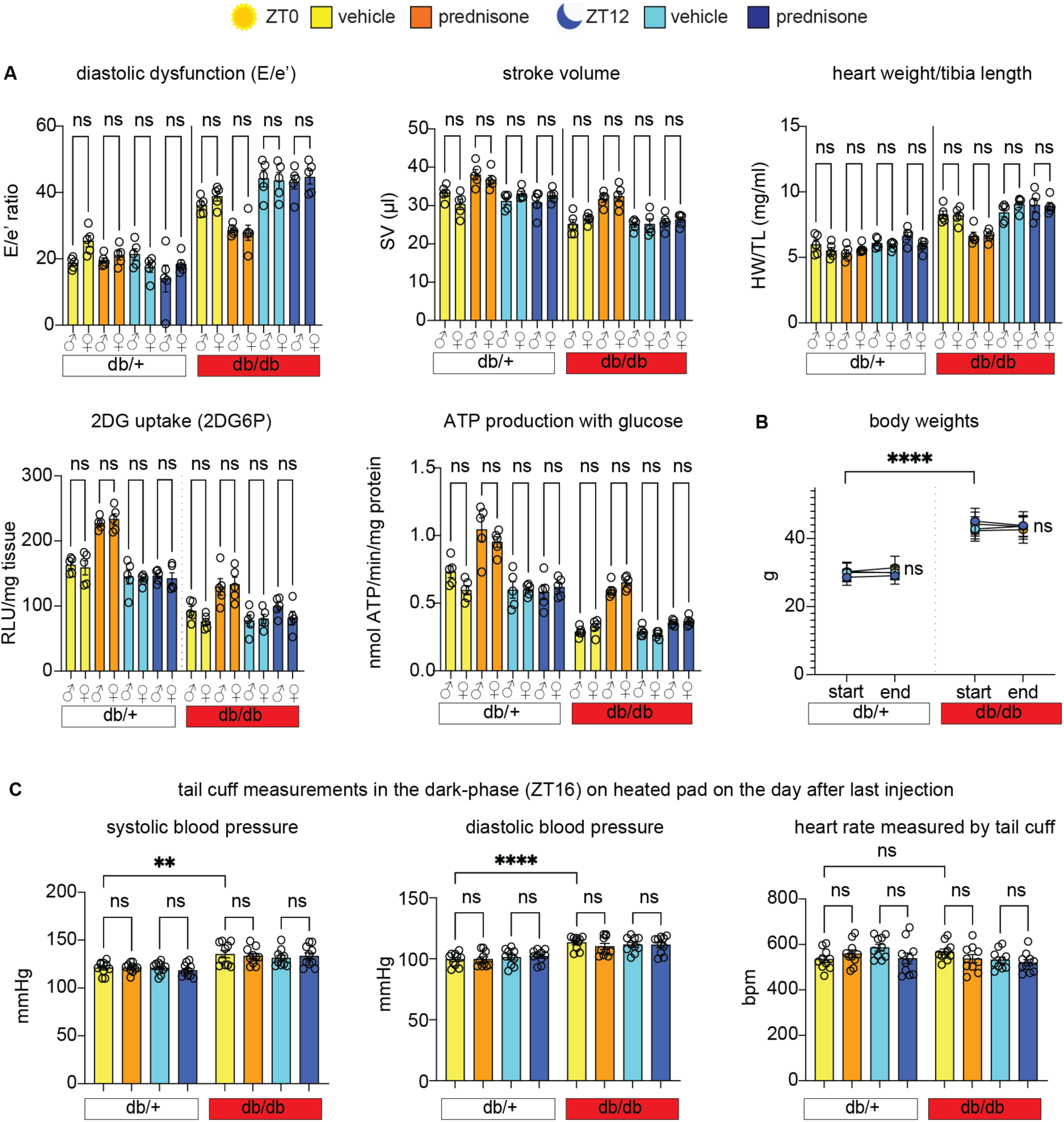
Additional measurements of cardiac function and blood pressure related to. Figure 5**. (A)** Sex-disaggregated measurements for male and female mice (db/+) and (db/db) for E/e’, stroke volume, heart weight/tibia length, 2DG uptake and ATP-linked respiration in cardiomyocytes showed no sizable sexual dimorphism in the treatment effects, either with ZT0 or ZT12 prednisone**. (B)** Treatments had no sizable effects on body weight, which was significantly skewed by genotype. **(C)** Treatments had no sizable effects on blood pressure or heart rate during blood pressure measurements beyond the expected db/db-related mild hypertension in both systole and diastole. Shown are mean±S.E.M, histograms show also individual mouse values. n=(5♀+5♂)/group; 3w ANOVA + Sidak; *, P<0.05; **, P<0.01; ***, P<0.001; ****, P<0.0001.

**Supplementary Figure 5.**
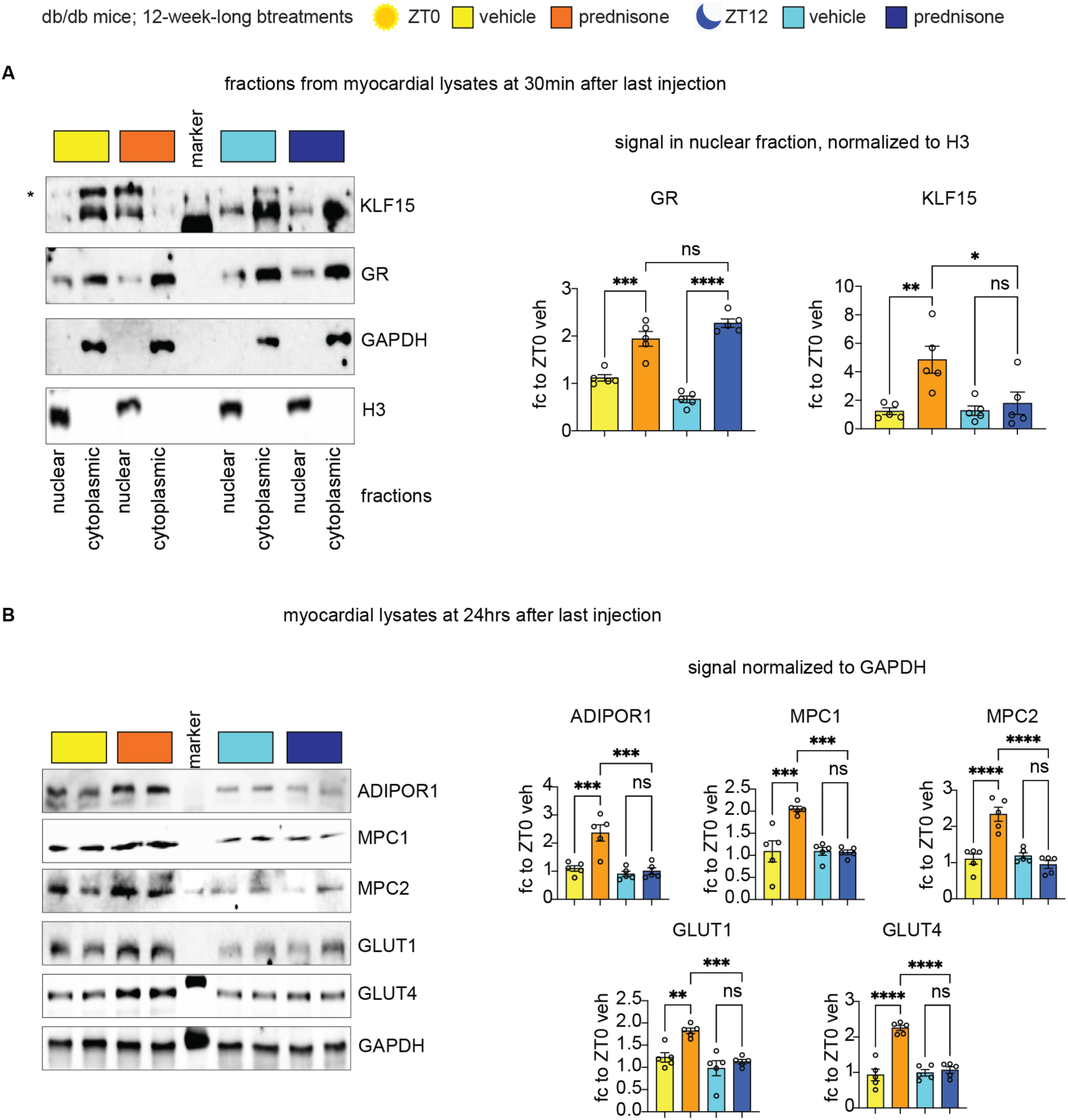
Additional protein analyses in db/db hearts related to. Figure 5**. (A)** WB in nuclear (H3^+^) and cytoplasmic (GAPDH^+^) fractions showed that time-of-intake changed treatment-driven nuclear translocation of KLF15 but not GR in db/db hearts. **(B)** ZT0 intermittent prednisone – but not ZT12 – increased the myocardial protein levels of Adipor1, Mpc1, Mpc2, Glut1 and Glut4 in db/db mice compared to vehicle. Shown are mean±S.E.M, histograms show also individual mouse values. n=5♂/group, 2w ANOVA + Sidak; *, P<0.05; **, P<0.01; ***, P<0.001; ****, P<0.0001.

## METHODS AND MATERIALS

### Sex as biological variable

We performed the bulk of our wild-type and diabetic mice experiments in both males and females, analyzing and reporting the physiological, molecular and histological assessments as sex-aggregated in main figures and sex-disaggregated in the supplementary figures. We did not find sizable sex-specific patterns. Therefore, we performed our unbiased omics-based screenings and subsequent KO- or knockdown-based proofs of requirement only in male mice to minimize variability and overall mouse number used.

### Animal handling and treatments

Mice were housed in a pathogen-free facility in accordance with the American Veterinary Medical Association (AVMA) and under protocols fully approved by the Institutional Animal Care and Use Committee (IACUC) at the Cincinnati Children’s Hospital Medical Center (#2023-0002). Consistent with the ethical approvals, all efforts were made to minimize suffering. Euthanasia was performed through carbon dioxide inhalation followed by cervical dislocation and heart removal.

Mice were maintained on a 12h/12h light/dark cycle, and diet/pharmacological treatments were initiated at ∼ 12 weeks of age. Mice were obtained and interbred from the Jakson Laboratories (Bar Harbor, ME; JAX strain number listed) or from collaborators: WT C57BL/6 mice #000664, *Myh6-MerCreMer* mice #005657, *GR-flox* mice #021021, *Arntl-flox* (BMAL1-flox) #007668, *Klf15-flox* mice #029787 (provided by Dr. Jain, Brown U), db/db #000642; Klf15-3xFLAG mice (25) were provided by Dr. Haldar (Amgen & UCSF). Gene ablation was induced with tamoxifen at ∼8 weeks of age i.p. tamoxifen (20 mg/kg/day for 5 days; Sigma #T5648) and analyses were performed after 4 weeks of tamoxifen washout, i.e. at ∼16 weeks of age.

Weekly prednisone treatment consisted of once-weekly intraperitoneal injection of prednisone (1 mg/kg; #P6254, Sigma-Aldrich, St. Louis, MO). The injectable solution was diluted from a stock (5 mg/ml) in dimethyl sulfoxide (DMSO; #D2650, Sigma-Aldrich, St. Louis, MO) in a 50-µl volume. Injections were conducted either at the beginning of the light phase (ZT0; lights on) or at the beginning of the dark phase (ZT12; lights off). Tissues were harvested at 4 hours post-injection for epigenetic/transcriptional assays and at 24 hours post-injection for metabolic/physiologic assays. All in vivo, ex vivo, and postmortem analyses were conducted blinded to treatment group.

For systemic AAV experiments, 6-month-old db/db mice were administered to either vehicle or 12-week long prednisone regimen at the beginning of the light phase (ZT0; lights on). These mice were further injected retro-orbitally with either 6×10^12^ genome copies/injection MyoAAV-scrambleshRNA or 1×10^12^ genome copies/injection for each of the MyoAAV-*Nr3c1*shRNA and MyoAAV-*Klf15*shRNA vectors (Vector Builder cargo vectors # VB010000-0023jze, VB240517-1300eyt, VB240517-1305cxu, VB240517-1306ywz, VB900144-1448eer, VB240517-1307ced, VB240517-1308zdv; anti-*Nr3c1* shRNA sequences: AGGATTGCAAGCCTCTTATTT, TGAGATTCGAATGACTTATAT, TTTGCTCCTGATCTGATTATT; anti-*Klf15* shRNA sequences: CATTTCTGCTTCCCTGAATTT, CTACCCTGGAGGAGATTGAAG, ACCGAAATGCTCAGTGGGTTA; U6 promoter for all scramble and shRNA constructs) while under inhaled isoflurane anesthesia. All MyoAAV injections were diluted in sterile PBS. To prepare and isolate AAV virions, we followed the procedures we previously reported (50).

### RNA-seq and qPCR analyses

RNA-seq was conducted on RNA extracted from left ventricle myocardial tissue. Each heart was immediately snap frozen in 1ml TRIsure (Bioline, BIO-38033) using liquid nitrogen. RNAs from each heart were re-purified using RNeasy Mini Kit (Cat #74104; Qiagen, Germantown, MD). RNA-seq was performed at the DNA Core (CCHMC). 150 ng – 300 ng of total RNA determined by Qubit (Cat #Q33238; Invitrogen, Waltham, MA) high-sensitivity spectrofluorometric measurement was poly-A selected and reverse transcribed using Illumina’s TruSeq stranded mRNA library preparation kit (Cat# 20020595; Illumina, San Diego, CA). Each sample was fitted with one of 96 adapters containing a different 8 base molecular barcode for high level multiplexing. After 15 cycles of PCR amplification, completed libraries were sequenced on an Illumina NovaSeqTM 6000, generating 20 million or more high quality 100 base long paired end reads per sample. A quality control check on the fastq files was performed using FastQC. Upon passing basic quality metrics, the reads were trimmed to remove adapters and low-quality reads using default parameters in Trimmomatic [Version 0.33]. In the next step, transcript/gene abundance was determined using kallisto [Version 0.43.1]. The trimmed reads were then mapped to mm10 reference genome using default parameters with strandness (R for single-end and RF for paired-end) option in Hisat2 [Version 2.0.5]. In the next step, transcript/gene abundance was determined using kallisto [Version 0.43.1]. We first created a transcriptome index in kallisto using Ensembl cDNA sequences for the reference genome. This index was then used to quantify transcript abundance in raw counts and counts per million (CPM). Differential expression (DE genes, FDR<0.05) was quantitated through DESeq2. PCA was conducted using ClustVis. Gene ontology pathway enrichment was conducted using the Gene Ontology analysis tool.

Total RNA was extracted from cryo-pulverized liver tissue and hiPSC-derived hepatocyte-like cells using Trizol (Thermo Fisher Scientific) and 1 ug RNA was reverse-transcribed using SuperScript^TM^ IV VILO^TM^ Master Mix (#11756050, Thermo Fisher Scientific). RT-qPCRs were conducted in three replicates using 1X SYBR Green Fast qPCR machine (Bio-Rad, Hercules, CA; thermal profile: 95C, 15 sec; 60C, 30sec; 40x; melting curve). The 2-ΔΔCT method was used to calculate relative gene expression. GAPDH was used as the internal control. Primers were selected among validated primer sets from MGH PrimerBank as follows (Gene, Forward, Reverse): *Klf15,* GAGACCTTCTCGTCACCGAAA, GCTGGAGACATCGCTGTCAT*; Adipor1,* TCTTCGGGATGTTCTTCCTGG, TTTGGAAAAAGTCCGAGAGACC*; Mpc1,* CTCCAGAGATTATCAGTGGGCG, GAGCTACTTCGTTTGTTACATGGC*; Mpc2,* CTCAGTCCACTGTGTTGATGGC, ATCCGAAACAGCTGAGAGGCTC.

### Mass-spec analysis of ceramides in myocardial tissue

Untargeted lipidomics was performed on myocardial tissue from mice treated for 12-week-long intermittent treatments with vehicle or weekly prednisone. Whole hearts were snap-frozen and then ground to powder with dry ice-chilled mortar and pestle. Frozen samples were sent on dry ice to mass spectrometry Core at the Cincinnati Children’s Hospital Medical Center, where LC-MS and data analysis were performed. Briefly 10 µl sample was added to 30 µl of internal standard, except for the blank calibrator. Then 300 µl of Butanol/Methanol (3:1) was added to sample aliquot tube and vortexed for 10 mins. Next, 150 µl of Heptane/Ethyl Acetate (3:1) was added to the sample tube and vortexed for 5 mins. The contents of the sample aliquot were then transferred to a large glass tube, to which 150 µl of Heptane/Ethyle Acetate was added and vortexed for 5 mins. Then, 300 ul of 1% Acetic acid was added to the original tube to wash, vortexed for 5 mins and then added back to the glass tube. The glass tube was further vortexed for 5 mins and let sit for 5 mins. The upper organic phase of ∼ 200 µl was transferred to a new glass tube. To this, 320 µl Heptane/Ethyl Acetate (3:1) was added to the water phase, vortexed for 5 mins and let sit for 5 mins. Next, 280 µl of upper organic phase (second extract) was transferred to the first extract. This step is repeated once. The upper organic phase (third extract) of 100 µl was combined with other extracts and dried down with nitrogen in a heating block set to 40°C. The resuspended lipid extracts (2 µl) were loaded and separated through Xevo QTOF G2-S mass spectrometer (Waters Corp). All calculations were performed using QuanLynx.

### Chromatin immunoprecipitation and sequencing

Whole hearts were cryopowdered using a liquid nitrogen cooled RETSCH Cryomill. The cryopowdered tissue was then fixed in 10 ml of 1% paraformaldehyde (PFA) for 30 min at room temperature with gentle nutation. Fixation was quenched 1 ml of 1.375 M glycine (Cat# BP381-5, Thermo Fisher Scientific, Waltham, MA) with gentle nutation for 5 mins at room temperature. After centrifugation at 3,000*g* for 5 mins at 4°C, the pellet was resuspended in cell lysis buffer as per reported conditions, supplementing the cell lysis buffer with cytochalasin B (3 µg/ml) and rotating for 10 min at 4°C. Nuclei were pelleted at 300*g* for 10 mins at 4°C and subsequently processed following the reported protocol with the adjustment of adding cytochalasin B (3 µg/ml) into all solutions for chromatin preparation and sonication, antibody incubation, and wash steps. Chromatin was then sonicated for 15 cycles (30 s, high powder, 30-s pause, and 500-µl volume) in a water bath sonicator set at 4°C (Bioruptor 300. Diagenode, Denville, NJ). After centrifuging at 10,000*g* for 10 mins at 4°C, sheared chromatin was checked on agarose gel for a shear band comprised between ∼150 and ∼600 bp. Two micrograms of chromatin were kept for pooled input controls, whereas ∼50 µg of chromatin was used for each pull-down reaction in a final volume of 2 ml rotating at 4°C overnight. Primary antibodies were as follows: rabbit poly-clonal anti-GR (#A2164, ABclonal), mouse monoclonal ANTI-FLAG M2 (#F1804, Sigma-Aldrich, St. Louis, MO). Chromatin complexes were precipitated with 100 µl of Dynabeads M-280 (Sheep anti-mouse, # 11202D; sheep anti-rabbit, #11204; Thermo Fisher Scientific, Waltham, MA). After washes and elution, samples were treated with proteinase K (Cat# 19131, QIAGEN, Hilden, Germany) at 55°C, and cross-linking was reversed through overnight incubation at 65°C. DNA was purified using a MinElute purification kit (Cat# 28004, QIAGEN, Hilden, Germany) and quantitated using Qubit reader and reagents. Library preparation and sequencing were conducted at the CCHMC DNA Core, using TruSeq ChIP-seq library prep (with size exclusion) on ∼10 ng of chromatin per ChIP sample or pooled inputs and HiSeq 50-bp was conducted using HOMER software (v4.10) after aligning fastq files to the mm10 mouse genome using bowtie2. PCA was conducted using ClustVis. Heatmaps of peak density were imaged with TreeView3. Peak tracks were imaged through WashU epigenome browser. Gene ontology pathway enrichment was conducted using the gene ontology analysis tool.

For ChIP-qPCR analyses, muscle chromatin was immunoprecipitated following the conditions for ChIP-seq. Input and IP chromatins were diluted 100X and assayed using the qPCR conditions. The regions identified in the Klf15 promoter were a GRE at −1167bp from TSS (primers, Fw TCTAGACAGCTGGGGCATCT, Rev GACAGACCTTCCTTCCTGGC) and a BMAL1 E-Box at −560bp from TSS (primers, Fw AAGCACAGACTCCTTCCGTG, Rev CGCTACCCTAGACTTCTGCG). Additional regions and primers: *Adipor1* GRE, Fw agctggggttctaggacact, Rev gctgctccccctttaagtgt; *Adipor1* KRE, Fw aagattgccttcccagctcc, Rev ggaggggccggaaatgttta; *Mpc1* GRE, Fw gctcttggttaaggcgacct, Rev acacatgattgtccggtccc; *Mpc1* KRE, Fw gctcttggttaaggcgacct, Rev acacatgattgtccggtccc; *Mpc2* KRE, Fw atagcgagatccaagccagc, Rev ctagggatcgacagcagcag; *Mpc2* GRE, Fw ctgctgctgtcgatccctag, Rev gctgagaccagacagacacc.Signal in IP chromatin was quantitated as % of input signal per qPCR calculations.

### 2-DG-6-phosphatase assay

For measuring the uptake of 2-DG-6-phosphatase, Glucose uptake-Glo^TM^ assay (Cat# J1341, Promega) was utilized to inject 1mM 2DG into mice for 30 mins before euthanasia. After 30 mins, the whole heart tissues were harvested and stored at −80C until ready to use. Next, the reagents from the kit were combined for reaction mixtures according to manufacturer’s instructions and incubated at room temperature for 1 hr. The frozen heart tissues were crushed into a fine powder and 20 - 50 mg of tissue powder was used for the assay. After 1 hour incubation of reaction mixture, 25 µl of Neutralization buffer was added to reaction mixture. Next, 125 µl of reaction mixture and neutralization buffer was added to powdered heart tissue and incubated for 1 hr. After incubation, the samples were centrifuged for 5 mins at 10,000*g* and 125 µl of the supernatant was loaded onto a 96-well plate for luminescence readings.

### Respirometry on cardiac biopsies, cardiomyocytes and isolated mitochondria

For cardiac biopsies, whole hearts were harvested and multiple transverse cuts were performed with a scapel in the left ventricular region with small scissors. Size matched tissue biopsies were prepared as thin as possible (∼0.5mm^2^) cover the bottom of the seahorse plate’s well. The biopsies were permeabilized in 100ug/ml saponin solution (Sigma #SAE0073) and incubated in XF base medium with 10 mM glucose, 2 mM L-glutamine, and 1 mM pyruvate (Sigma #G7021, #G7513, #P2256) in 37°C non-CO_2_ incubator for 30 mins. Regular seahorse protocol is followed with basal reads and injections of oligomycin, FCCP, Rotenone/Antimycin following reported conditions. Basal OCR was calculated as baseline value (average of 3 consecutive reads) minus value after rotenone/antimycin addition (average of 3 consecutive reads). Basal OCR values were normalized to total protein content using Bradford assay.

For cardiomyocytes, respirometry was performed on freshly isolated cells. Cardiomyocytes are isolated adapting to previously reported conditions. In anesthetized mice, the descending aorta is cut, and the heart is immediately flushed by injecting 7ml pre-warmed EDTA buffer into the right ventricle. Immediately afterwards, the ascending aorta is clamped using Reynolds forceps, the heart is removed and kept in the 60 mm dish with pre-warmed EDTA buffer. The digestion process is proceeded by injecting 10 ml pre-warmed EDTA buffer to the apex of left ventricle. The heart is then transferred to the 60 mm dish with perfusion buffer and 3 ml perfusion buffer is injected to flush out the EDTA buffer. Then the heart is transferred to another 60 mm dish with pre-warmed 180U/ml Collagenase II buffer and 30-40 ml collagenase buffer is injected into the apex of LV. Then the heart is cut from the clamp, the ventricular myocardial tissue is pulled gently into small pieces using forceps and further dissociated by gentle pipetting. The cardiomyocyte suspension is passed through a 200 µm mesh, and the digestion is stopped by adding 5ml of stop buffer. Cardiomyocytes are then centrifuged at 100*g* for 3 mins, resuspended in the stop buffer with increasing concentration of calcium (100 µM, 400 µM, and 1000 µM), re-centrifuged and then plated in laminin-coated Seahorse plates in culture media. After 1 h incubation, the culture media is aspirated and Seahorse media is added for respirometry, which starts after an additional 1 h equilibrium in a CO_2_-free incubator. Buffer compositions: EDTA buffer contains 130 mM NaCl, 5mM KCl, 0.5mM NaH_2_PO_4_, 10mM HEPES, 10mM glucose, 10mM BDM, 10mM Taurine, 5 mM EDTA, pH 7.8; Perfusion buffer contains 130 mM NaCl, 5 mM KCl, 0.5 mM NaH_2_PO_4_, 10 mM HEPES, 10 mM glucose, 10 mM BDM, 10 mM taurine, 1 mM MgCl_2_, pH 7.8; Collagenase buffer contains 220U/ml Collagenase II; Stop buffer consists of Perfusion buffer supplemented with 5% sterile fetal bovine serum; Culture medium (250 mL) contains 2.45g Hanks’ salt, 5 mL non-essential amino acids, 2.5 mL MEM Vitamin Solution, 0.0875g NaHCO_3_, 2.5 mL PenStrep 10X, 1 mg/mL bovine serum albumin. Regular seahorse protocol is followed with basal reads and injection of oligomycin, FCCP, Rotenone/Antimycin following reported conditions. Basal OCR was calculated as baseline value (average of 3 consecutive reads) minus value after rotenone/antimycin addition (average of 3 consecutive reads). Basal OCR values were normalized to total protein content, assayed in each well after the Seahorse ran through homogenization and Bradford assay.

Mitochondrial ATP production rate was calculated as OCR_ATP_ ∗ 2 (mol O in mol O_2_) ∗ 2.75 (Seahorse P/O), where OCR_ATP_ is the difference between baseline and oligomycin OCR. Nutrients: 10 mM glucose or 10 mM pyruvate; inhibitors: 0.5 μM rotenone + 0.5 μM antimycin A (Agilent). Respiratory control ratio (RCR) values were obtained from isolated mitochondria from myocardial tissue. Tissues from the left ventricle are harvested from the mouse and cut up into very fine pieces. The minced tissue is placed in a 15 mL conical tube (USA Scientific #188261) and 5 mL of MS-EGTA buffer with 1 mg Trypsin (Sigma #T1426-50 MG) is added to the tube. The tube is quickly vortexed, and the tissue is left submerged in the solution. After 2 min, 5 mL of MS-EGTA buffer with 0.2% BSA (Goldbio #A-421-250) is added to the tube to stop the trypsin reaction. MS-EGTA buffer: Mannitol (#M0214-45, ChemProducts), Sucrose (#100892, Millipore), HEPES (#15630–080, Gibco), EGTA (#E14100–50.0, RPI). The tube is inverted several times to mix then set to rest. Once the tissue has settled to the bottom of the tube, 3 mL of buffer is aspirated, and the remaining solution and tissue is transferred to a 10 mL glass tissue homogenizer (Avantor # 89,026–382). Once sufficiently homogenized the solution is transferred back into the 15 mL conical tube and spun in the centrifuge at 1,000g for 5 min at 4 °C. After spinning, the supernatant is transferred to a new 15 mL conical tube. The supernatant in the new tube is then centrifuged at 12,000g for 10 min at 4 °C to pellet the mitochondria. The supernatant is discarded from the pellet and the pellet is then resuspended in 7 mL of MS-EGTA buffer and centrifuged again at 12,000g for 10 min at 4 °C. After spinning, the supernatant is discarded, and the mitochondria are resuspended in 1 mL of Seahorse medium (Agilent #103335–100) with supplemented 5 mM pyruvate (Sigma #P2256-100G). After protein quantitation using a Bradford assay (Bio-Rad #5000001), 2.5 μg mitochondria are dispensed per well in 180 μl total volumes and let to equilibrate for 1 h at 37 degrees C. 20 μl of 5 mM ADP (Sigma #01905), 50 μM Oligomycin (Milipore #495455-10 MG), 100 μM Carbonyl cyanide-p-trifluoromethoxyphenylhydrazone (TCI #C3463), and 5 μM Rotenone (Milipore #557368-1 GM)/Antimycin A (Sigma #A674-50 MG) are added to drug ports A, B, and C, respectively. At baseline and after each drug injection, samples are read three consecutive times. RCR was calculated as the ratio between state III (OCR after ADP addition) and uncoupled state IV (OCR after oligomycin addition). All Seahorse measurements were conducted blinded to treatment groups.

### Western blotting

Protein analyses in liver were performed on ∼ 50 ug total lysates from the whole heart. Cyro-pulverized liver tissue was incubated in RIPA buffer (Cat #89900, Thermo Fisher Scientific, Waltham, MA) supplemented with 1x protease/phosphatase inhibitor (Cat #78440, Thermo Fisher Scientific, Waltham, MA) for 30 mins and sonicated for 10 secs twice. The samples were then centrifuged at 10,000 rpm for 10 mins at 4°C. Supernatant containing the protein is transferred into a new tube and used as a total lysate. For total cell lysates from culture cells, cells were harvested and resuspended in RIPA buffer containing 1x protease and phosphatase inhibitors. Lysates were incubated for 30 mins and centrifuged at 10,000 rpm for 10 mins at 4°C. The supernatant was used as a total cell lysate. The protein concentrations of the supernatants were determined using the Pierce BCA Protein Assay kit (Cat #23227, Thermo Fisher Scientific, Waltham, MA). Equal amounts of protein were separated using SDS-PAGE and transferred to a PVDF membrane (Cat #1620174, BioRad, Hercules, CA). Membranes were blocked in 5% milk in TBST for 1 hour at room temperature and then incubated overnight at 4°C with primary antibodies: Mouse Klf15 monoclonal antibody (Cat# sc-271675, Santa Cruz Biotechnology), Rabbit Gapdh monoclonal antibody (Cat# A10868, AC001 ABclonal), Rabbit Histone H3 polyclonal antibody (Cat# A2348, ABclonal), Mouse GR monoclonal antibody (Cat# sc-393232, Santa Cruz Biotechnology), Rabbit AdipoR1 polyclonal antibody (Cat# A16527, ABclonal), Rabbit Mpc1 polyclonal antibody (Cat# A20195, ABclonal), Rabbit Mpc2 polyclonal antibody (Cat# A20196, ABclonal), Rabbit Glut1 polyclonal antibody (Cat# A11208, ABclonal), Rabbit Glut4 polyclonal antibody (Cat# A7637, ABclonal) followed by incubation with HRP-conjugated secondary antibodies: Peroxidase AffiniPure Goat Anti-Rabbit IgG (H+L) (Cat# 111-035-003, Jackson ImmunoResearch Labs, West Grove, PA) for 1 hour at room temperature. Immunoreactive bands were visualized by chemiluminescence using Pierce Enhanced Chemiluminescent western blotting substrate (Cat #32106, Thermo Fisher Scientific, Waltham, MA).

### Nuclear and cytoplasmic fraction analysis

The separation of nuclear and cytoplasmic protein analysis was performed using NE-PER nuclear and cytoplasmic extraction kit (Invitrogen #78835). Briefly, 100 mg of the heart tissues were homogenized, and 1 ml of CERI solution was added and vortexed vigorously on high setting for 15 sec. Following incubation on ice for 10 mins, 55 µl od ice-cold CERII solution was added, vortexed, and incubated for 1 minute and centrifuged at 16,000g for 5 mins. The supernatant containing cytoplasmic fraction was separated and collected into a new 1.5 ml Eppendorf tube. The insoluble is suspended in 500 µl of ice-cold NER solution. The sample was then placed on ice for 40 mins and vortexed every 10 mins for 1 sec. Finally, the samples were centrifuged at 16,000g for 10 mins. The supernatant containing the nuclear fraction was separated into a new 1.5 ml Eppendorf tube. All samples were stored at −80°C until use. The protein concentration was determined by Bio-Rad protein assay as described previously. Once the protein concentration was determined, western blotting analysis was carried out as previously described.

### Measurement of Blood pressure by Tail-cuff Plethysmography

The CODA 8 system (Kent Scientific, Torrington, CT) was used to noninvasively measure blood pressure in mice through tail-cuff recordings. It was factory calibrated, and standard settings and procedures were adhered to as recommended by the manufacturer. Before beginning the experiments, the occlusion and VPR cuffs were routinely checked. Experiments were carried out in a controlled environment maintained at 22 ± 2°C, where mice were acclimated for an hour prior to measurements. Mice were gently guided into the restraint tubes and tube end holders were adjusted at the tail’s base, with the VPR sensor cuff positioned next to it. Heating pads provided with the CODA 8 system were preheated to 33-35°C. Mice were warmed on these pads for 5 mins before and during the blood pressure recording. To obtain blood pressure readings, the occlusion cuff was inflated to 25 mm Hg and then deflated over 20 seconds. The VPR sensor cuff detected the changes in tail volume as blood flow returned deflation, with a minimum volume change set at 15 µl. Each recording session included 15-25 cycles of inflation and deflation, with the first 5 cycles used for acclimation and excluded from the analysis. The remaining cycles were used for data collection. Mice were habituated to the procedure for at least 3 days before blood pressure measurements were recorded.

### Echocardiography

Mice were anesthetized with 1.5% isoflurane (1.5% oxygen) and imaged in the supine position using Vevo 3100 Imaging System with a 35-MHz linear probe (Visualsonics, Canada). The core temperature was maintained at 37°C. Heart rates were kept consistent between the groups (400 – 450 bpm) (51). The heart was imaged in a 2D, short axis, and 4-chamber view by placing the pulsed wave Doppler at the septal corner of the mitral annulus. Mitral inflow velocity (E) and longitudinal tissue velocity of the mitral anterior annulus (e’) were measured through Vevo2100 1.5.0 Image Software (52).

### Statistics

Statistical analyses were performed using Prism software v8.4.1 (Graphpad, La Jolla, CA). The Pearson-D’Agostino normality test was used to assess data distribution normality. When comparing two groups, two-tailed Student’s t-test with Welch’s correction (unequal variances) was used. When comparing data for two or three interacting variables, two-way and three-way ANOVA tests were used respectively. P value less than 0.05 was considered significant. When the number of data points was less than 10, data were presented as single values (dot plots, histograms). Analyses pooling data points over time were presented as line plots connecting means and standard error of the mean.

### Study approval

Mice were housed in a pathogen-free facility in accordance with the American Veterinary Medical Association (AVMA) and under protocols fully approved by the Institutional Animal Care and Use Committee (IACUC) at Cincinnati Children’s Hospital Medical Center (#2022-0020, #2023-0002).

### Data availability

RNA-seq and ChIP-seq datasets reported here are available on GEO as GSE252826, GSE252857, GSE252856, GSE254114.

## Author contributions

HBD, ADP, FEAS, HBD, HL, KMF, KM, KP, CW: Data curation, Formal analysis, Investigation; MKJ, SMH: Resources; MQ: Conceptualization, Formal analysis, Funding acquisition, Supervision. Order of co-first Authors was assigned based on overall weight of contribution considering initial and revised submissions.

